# Customizing the Structure of Minimal TIM Barrels to Craft Efficient *De Novo* Enzymes

**DOI:** 10.1101/2025.01.28.635154

**Authors:** Julian Beck, Benjamin J. Smith, Mark Kriegel, Niayesh Zarifi, Emily Freund, Ahana G. Harsha, Jan Hartmann, Roberto A. Chica, Birte Höcker

**Author notes:** These authors contributed equally (Julian Beck, Benjamin J. Smith). **Corresponding authors:** Birte Höcker, Roberto A. Chica.

## Abstract

The TIM barrel is the most prevalent fold in natural enzymes, supporting efficient catalysis of diverse chemical reactions. While *de novo* TIM barrels have been successfully designed, their minimalistic architecture lacks structural elements essential for substrate binding and catalysis. Here, we present CANVAS, a computational workflow that introduces a structural lid into a minimal *de novo* TIM barrel to anchor catalytic residues and form an active-site pocket for enzymatic function. Starting from two *de novo* TIM barrels, we designed nine variants with distinct lids to form active sites for the Kemp elimination. Four designs showed measurable activity, with the most active reaching a catalytic efficiency of 21,000 M⁻¹ s⁻¹ at its optimal pH. A co-crystal structure of this variant bound to a transition-state analogue confirmed the accuracy of the designed lid and active site. Using the X-ray structure of a lower-activity variant (19 M⁻¹ s⁻¹), we applied ensemble-based design to optimize its active site, increasing catalytic efficiency by >1,600-fold to 32,000 M⁻¹ s⁻¹. These results demonstrate that *de novo* TIM barrels can be endowed with substrate binding pockets supporting efficient catalytic function, establishing a platform for building enzymes on demand from minimal protein scaffolds.

---

The TIM barrel is one of the most versatile enzyme folds. It is present in six of the seven enzyme classes and supports catalysis at diffusion-limited rates^1–3^. For these reasons TIM barrels have frequently been used as scaffolds for enzyme design^4–7^. While idealized *de novo* TIM barrels have been successfully designed^8–11^, none display enzymatic function due to their minimalistic architecture, which lacks key structural elements required for catalysis, such as cavities, pockets, and extended loops. Efforts to address these deficiencies have led to the introduction of secondary structural elements to create rudimentary pockets^12–14^. However, these pockets are too solvent exposed and undefined to support the microenvironments required for catalysis. *De novo* TIM barrels have also been fused to additional protein domains to create hybrid architectures with metal-binding sites positioned at domain interfaces^15^. While these constructs demonstrate that domain fusion can generate large cavities acting as reaction chambers for photoredox catalysis^16^, they rely largely on the inherent reactivity of the bound metal and lack substrate-specific active sites characteristic of natural TIM barrel enzymes. Therefore, novel strategies are needed to craft custom active sites and endow *de novo* TIM barrels with true enzymatic function.

## A computational approach for active-site scaffolding

Here, we introduce CANVAS (Customizing Amino-acid Networks for Virtual Active-site Scaffolding), a computational workflow for transforming minimal proteins such as *de novo* TIM barrels into functional enzymes (Figure 1a, Supplementary Figure 1). The process begins with a minimal TIM-barrel template and a theozyme, which is a computational model of an idealized enzyme active site with catalytic groups arranged to stabilize a reaction’s transition state^17^. Using the protein design software Triad^18^, a catalytic amino acid from the theozyme is placed onto the TIM-barrel template, and the transition state is built from its side chain to maintain the catalytic geometry^19^. A second catalytic residue is then built from the transition state, ensuring proper geometry for catalytic interaction. The C_α_ of this second residue is thus positioned in the empty space above the TIM barrel catalytic face. This allows it to serve as an anchor for constructing a custom lid to form the active-site pocket, which is designed using the generative AI tool RFdiffusion^20^ followed by sequence optimization with ProteinMPNN^21^ and structure prediction with AlphaFold2^22^. The active site is then optimized with Triad to build a pocket complementary to the transition state^6^. Designs are filtered using key enzyme design criteria^23^, including solvent-accessible surface area of the transition state, active-site preorganization, energy, and catalytic contact geometry.

**Figure 1.**
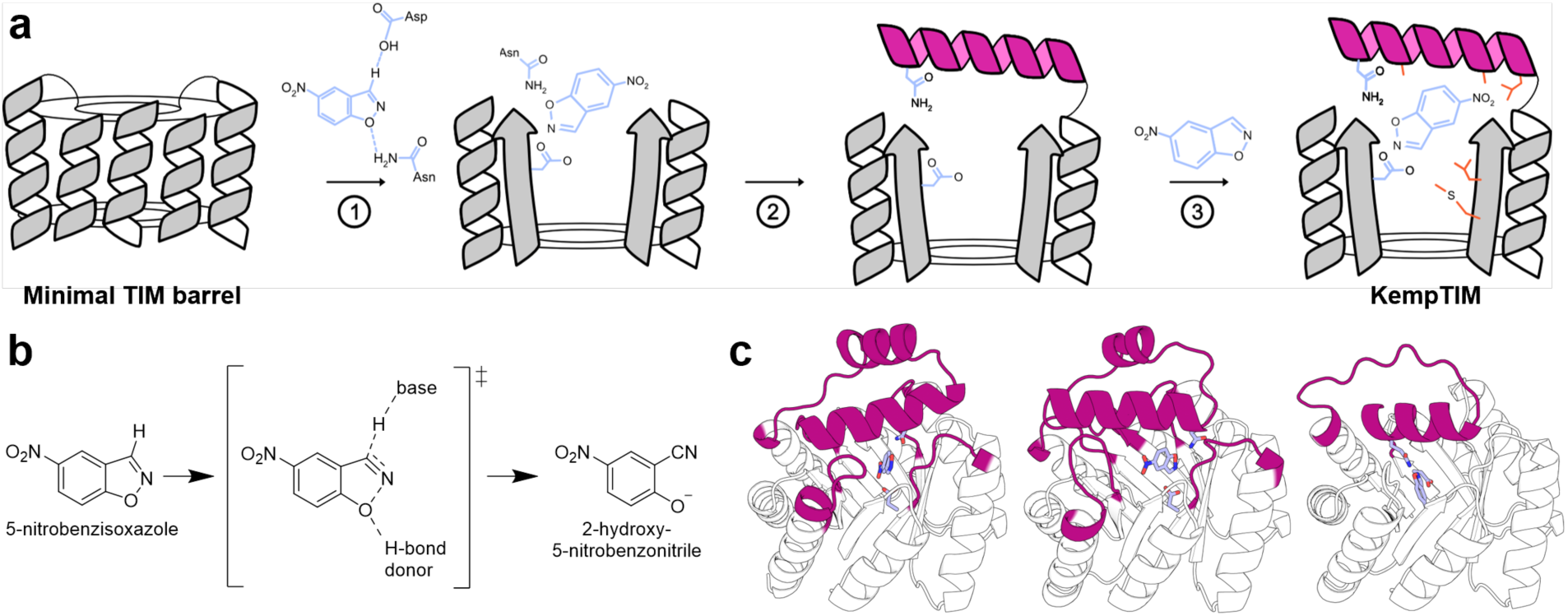
*De novo* enzyme design using CANVAS. (a) CANVAS follows three general steps: (1) placement of the theozyme (blue) onto the minimal TIM-barrel scaffold using a single catalytic residue as an anchor; (2) construction of a custom lid (magenta) to form the active-site pocket and anchor the second catalytic residue; and (3) active-site repacking to further stabilize the transition state. (b) Kemp elimination reaction. (c) Structure of three distinct lids (magenta) tailored to the Kemp elimination, designed using CANVAS. Transition state and designed catalytic residues are shown as sticks.

## Applying CANVAS to minimal TIM-barrel proteins

We applied CANVAS to the Kemp elimination (Figure 1b), a model organic transformation commonly used as a benchmark for *de novo* enzyme design^4,6,23^. The reaction’s theozyme features an Asp as catalytic base and an Asn or Gln as H-bond donor to stabilize the binding of the substrate and properly orient it for reaction with the base. We used the crystal structures of two *de novo* TIM barrels (PDB IDs: 7MCD^24^ and 7OSV^9^) as templates. Since 7OSV has its N-and C-termini on the catalytic face of the TIM barrel, we applied an *in silico* circular permutation based on the inpainting method^25^ to generate two modified structures termed NT6-CP1 and NT6-CP2 that contain termini on the stability face (Supplementary Figure 2). Starting from these structures, we used CANVAS to generate distinct lids for each template tailored to the Kemp elimination (Figure 1c). The lid diffused onto 7MCD consists of a single 26-residue fragment introduced in βα-loop 7 (Supplementary Figure 3). By contrast, the designed lids in NT6-CP1 and NT6-CP2 comprise a major fragment of 33 or 39 residues introduced into βα-loops 2 or 6, respectively, to scaffold the catalytic H-bond donor, as well as three or four elongated loops to enhance interactions with the primary fragment. The resulting designs include variants 1–2 obtained from NT6-CP2, variants 3–5 from NT6-CP1 and variants 6–9 from 7MCD, each with a unique active-site configuration (Supplementary Figure 4) and sequence (Supplementary Table 1), which yielded high AlphaFold2 pLDDT scores (Supplementary Figure 5).

## Several active Kemp eliminases identified

Of the nine designed variants, those derived from NT6-CP1 and NT6-CP2 (variants 1–5) were either non-expressible or poorly soluble, whereas 7MCD-derived constructs (variants 6–9) were readily expressed and soluble. Of these initial designs, only variant 4 showed detectable activity; however, detailed characterization was hampered by low protein yields and a tendency to aggregate (Supplementary Figure 6, Supplementary Table 2). The NT6-CP1 template itself also failed to express, in contrast to NT6-CP2 and 7MCD. To improve expression and solubility of variants 1–5, we used LigandMPNN to redesign all non-active-site residues while preserving the Triad-designed active site. These new sequences are henceforth termed KempTIMs (Figure 2a, Supplementary Figure 7a, Supplementary Table 1). This strategy, previously applied to stabilize natural enzymes^26^, restored expression or increased solubility in all but one variant. The four soluble KempTIMs were purified at yields of 7–59 mg L⁻¹ (Supplementary Table 2) as monomeric species (Figure 2b, Supplementary Figure 7b, Supplementary Table 3) with mixed α/β secondary structure characteristic of TIM barrels (Figure 2c, Supplementary Figure 7c). All displayed melting temperatures above 65 °C, and several remained folded at temperatures exceeding 90 °C (Figure 2d, Supplementary Figure 7d).

**Figure 2.**
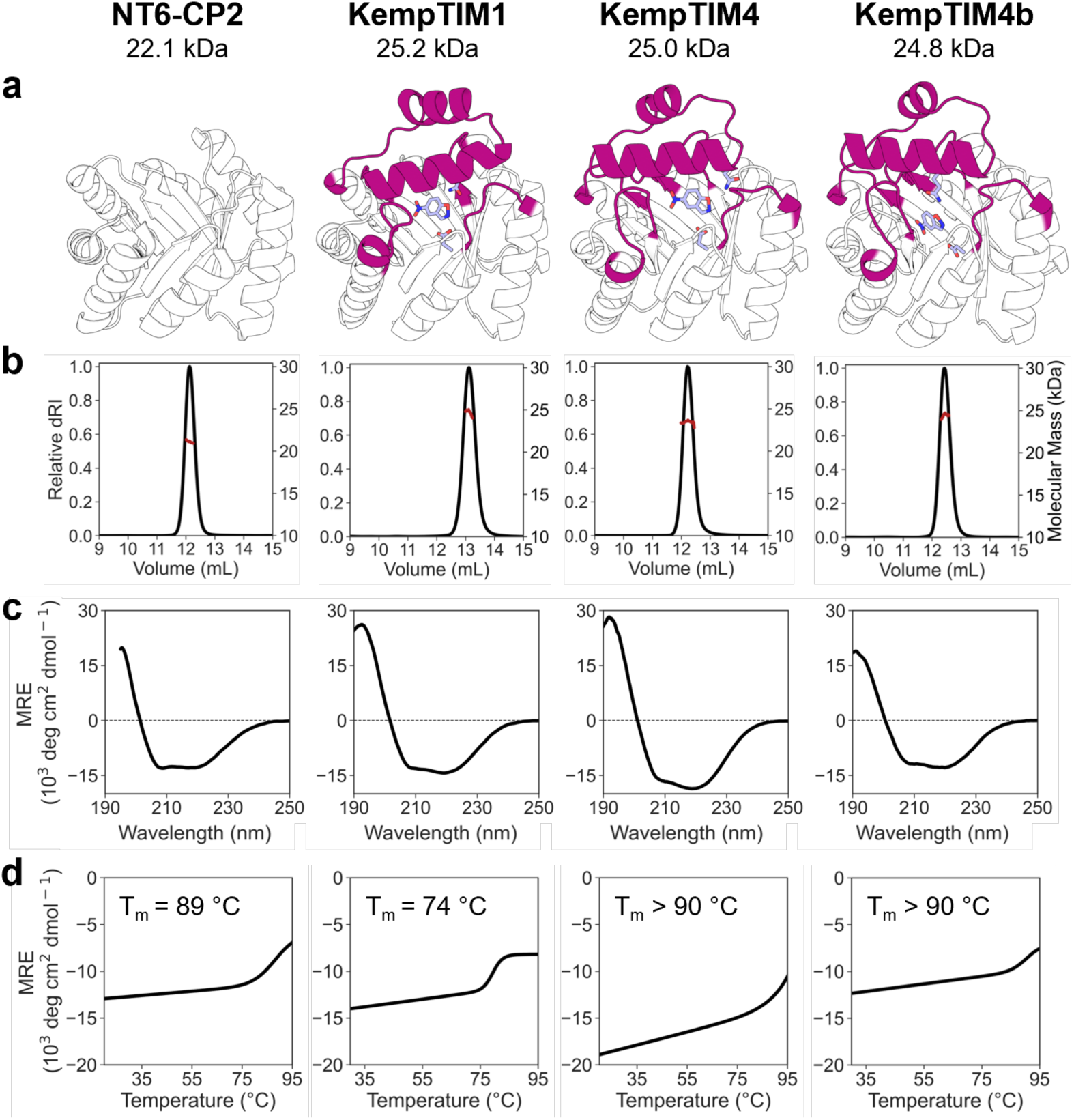
Structural characterization. (a) Computational models of minimal TIM-barrel NT6-CP2 and KempTIMs with the designed lid shown in magenta and the theozyme as sticks. (b) SEC-MALS indicates that the proteins are monomeric, with molecular weights close to their theoretical values. dRI: differential refractive index. (c) Circular dichroism spectra reveal a mixed αβ signal characteristic of TIM barrels. MRE: mean residue ellipticity. (d) Melting curves demonstrate that KempTIM1 is substantially destabilized compared to NT6-CP2. T_m_: melting temperature.

Kinetic analysis revealed activity for all expressed KempTIMs (Figure 3a, Supplementary Figure 8, Supplementary Table 4). The most active design, KempTIM1, exhibited a catalytic efficiency of 1,400 M^−1^ s^−1^ at pH 7 (Table 1). Mutation of the catalytic base abolished activity (Figure 3b), whereas mutation of the H-bond donor decreased *k*_cat_ sixfold (Figure 3c, Table 1), without disrupting the structure (Supplementary Figure 9), confirming the intended functional roles of the designed residues. A pH-rate profile (Supplementary Figure 10, Supplementary Table 5) revealed increasing activity at alkaline pH, with a single p*K*_a_ of approximately 9.0, consistent with the catalytic base Asp residing in a hydrophobic active-site environment. At its optimal pH of 10, *k*_cat_/*K*_M_ increased to 21,000 M^−1^ s^−1^ (Figure 3d, Table 1). KempTIM1 displayed *k*_cat_ values of 4.2 s^−1^ at pH 7.0 and 13.7 s^−1^ at pH 10.0, representing 10–14-fold increases compared to recently reported first-round Kemp eliminases assayed under similar conditions^27,28^. Together, these results show that CANVAS reliably produces functional *de novo* enzymes and that diverse lid architectures with distinct catalytic motifs can support Kemp elimination. Notably, the top-performing KempTIM surpasses the catalytic efficiencies of first-round *de novo* Kemp eliminases produced through alternative computational design strategies^27,28^.

**Figure 3.**
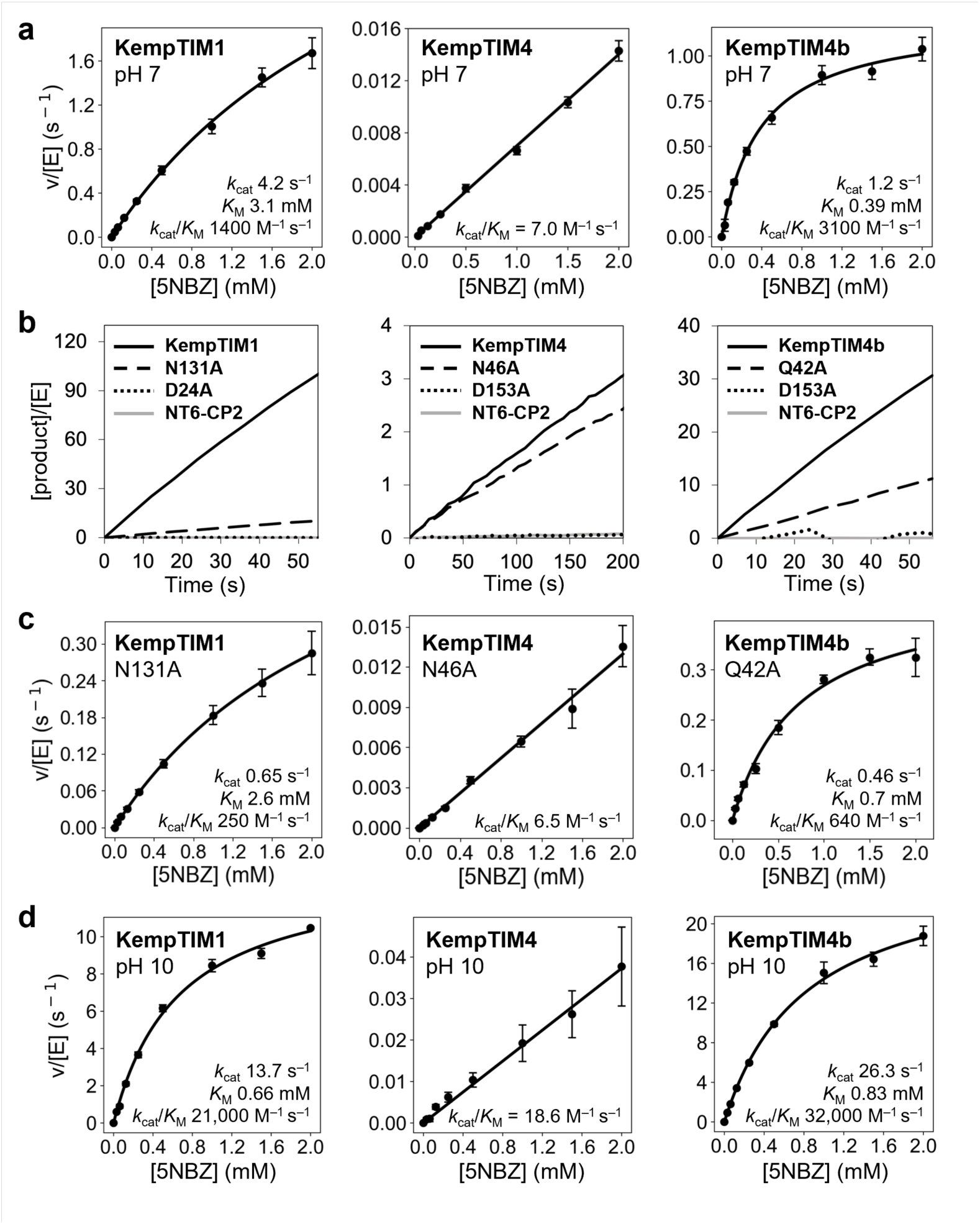
Kinetic characterization. Assays were carried out in 50 mM sodium phosphate containing 100 mM NaCl and 10% MeOH (pH 7) or 50 mM CHES containing 100 mM NaCl and 10% MeOH (pH 10). (a) Michaelis-Menten plots (pH 7) showing normalized initial rates as a function of 5-nitrobenzisoxazole (5NBZ) concentration. Data represent the mean ± SEM of ≥ 5 replicate measurements from ≥ 2 independent protein batches. (b) Reaction progress curves (pH 7) at 2 mM 5NBZ illustrating the contributions of the designed catalytic residues to overall catalysis. (c) Michaelis-Menten plots (pH 7) for variants lacking H-bond donor residues. Data represent the mean ± s.d. of ≥ 3 replicate measurements from one independent protein batch. (d) Michaelis-Menten plots (pH 10) showing normalized initial rates as a function of 5NBZ concentration. Data represent the mean ± SEM of ≥ 4 replicate measurements from ≥ 2 independent protein batches.

**Table 1.** Kinetic parameters of selected KempTIMs.

| Enzyme | Base | H-bond donor | pH | $k_{\text{cat}}$<br>(s<sup>-1</sup>) | $K_M$<br>(mM) | $k_{\text{cat}}/K_M$<br>(M<sup>-1</sup> s<sup>-1</sup>) |
| --- | --- | --- | --- | --- | --- | --- |
| KempTIM1 | D24 | N131 | 7 | 4.2 ± 0.2 | 3.1 ± 0.2 | 1,400 ± 100 |
|  |  |  | 10 | 13.7 ± 0.5 | 0.66 ± 0.06 | 21,000 ± 2,000 |
|  | D24A | N131 | 7 | N.D. | N.D. | N.D. |
|  | D24 | N131A | 7 | 0.65 ± 0.02 | 2.6 ± 0.1 | 250 ± 10 |
| KempTIM4 | D153 | N46 | 7 | N.D. | N.D. | 7.01 ± 0.08 |
|  |  |  | 10 | N.D. | N.D. | 18.6 ± 0.4 |
|  | D153A | N46 | 7 | N.D. | N.D. | N.D. |
|  | D153 | N46A | 7 | N.D. | N.D. | 6.5 ± 0.1 |
| KempTIM4b | D153 | Q42 | 7 | 1.21 ± 0.04 | 0.39 ± 0.04 | 3,100 ± 300 |
|  |  |  | 10 | 26.3 ± 0.9 | 0.83 ± 0.06 | 32,000 ± 3,000 |
|  | D153A | Q42 | 7 | N.D. | N.D. | N.D. |
|  | D153 | Q42A | 7 | 0.46 ± 0.03 | 0.7 ± 0.1 | 640 ± 90 |

$k_{\text{cat}}$ and $K_M$ using 5-nitrobenzisoazole as substrate were calculated by fitting the data to the Michaelis-Menten model $v_0 =$ $k_{\text{cat}}[E_0][S]/(K_M + [S])$ . Errors of nonlinear regression fitting, which represent the absolute measure of the typical distance that each data point falls from the regression line, are provided. N.D. indicates that no activity was detected, or that $K_M$ and $k_{\text{cat}}$ could not be determined accurately because saturation was not possible at the maximum concentration of 5-nitrobenzisoazole tested (2 mM), which is the substrate's solubility limit. In those cases, catalytic efficiencies ( $k_{\text{cat}}/K_M$ ) were calculated from the slope of the linear portion ( $[S] \ll K_M$ ) of the Michaelis-Menten model ( $v = (k_{\text{cat}}/K_M)[E_0][S]$ ). n = 2 independent protein batches for KempTIM1 and KempTIM4b. n = 4 independent protein batches for KempTIM4. At least two technical replicates were performed for each independent batch. Kinetic assays were carried out in 50 mM sodium phosphate containing 100 mM NaCl and 10% MeOH (pH 7) or 50 mM CHES containing 100 mM NaCl and 10% MeOH (pH 10).

## Confirming the lid and active-site features

To assess structural accuracy of the CANVAS-designed lids and their associated active sites, we solved crystal structures (Supplementary Table 6) of the most efficient design, KempTIM1 (21,000 M^−1^ s^−1^), and a lower-efficiency variant incorporating a different lid, KempTIM4 (19 M^−1^ s^−1^). In both enzymes, the designed lids fold as intended, with the helix–turn–helix motif and elongated loops closely matching the design models (lid backbone Cα RMSD of 0.9 Å for KempTIM1 and 1.2 Å for KempTIM4; Figure 4a,c). In both cases, however, the lid is slightly displaced relative to the TIM-barrel core, shifting the Cα position of the catalytic H-bond donor by 1.0 Å in KempTIM1 and 2.0 Å in KempTIM4 relative to the design models (Figure 4b,d). Because extensive crystal contacts were observed in the lid region of KempTIM4, we probed its solution structure using small-angle X-ray scattering (SAXS). Comparison of different structural models with the SAXS data indicates that the lid adopts a conformation in solution similar to that observed in the crystal structure (Supplementary Figure 11).

**Figure 4.**
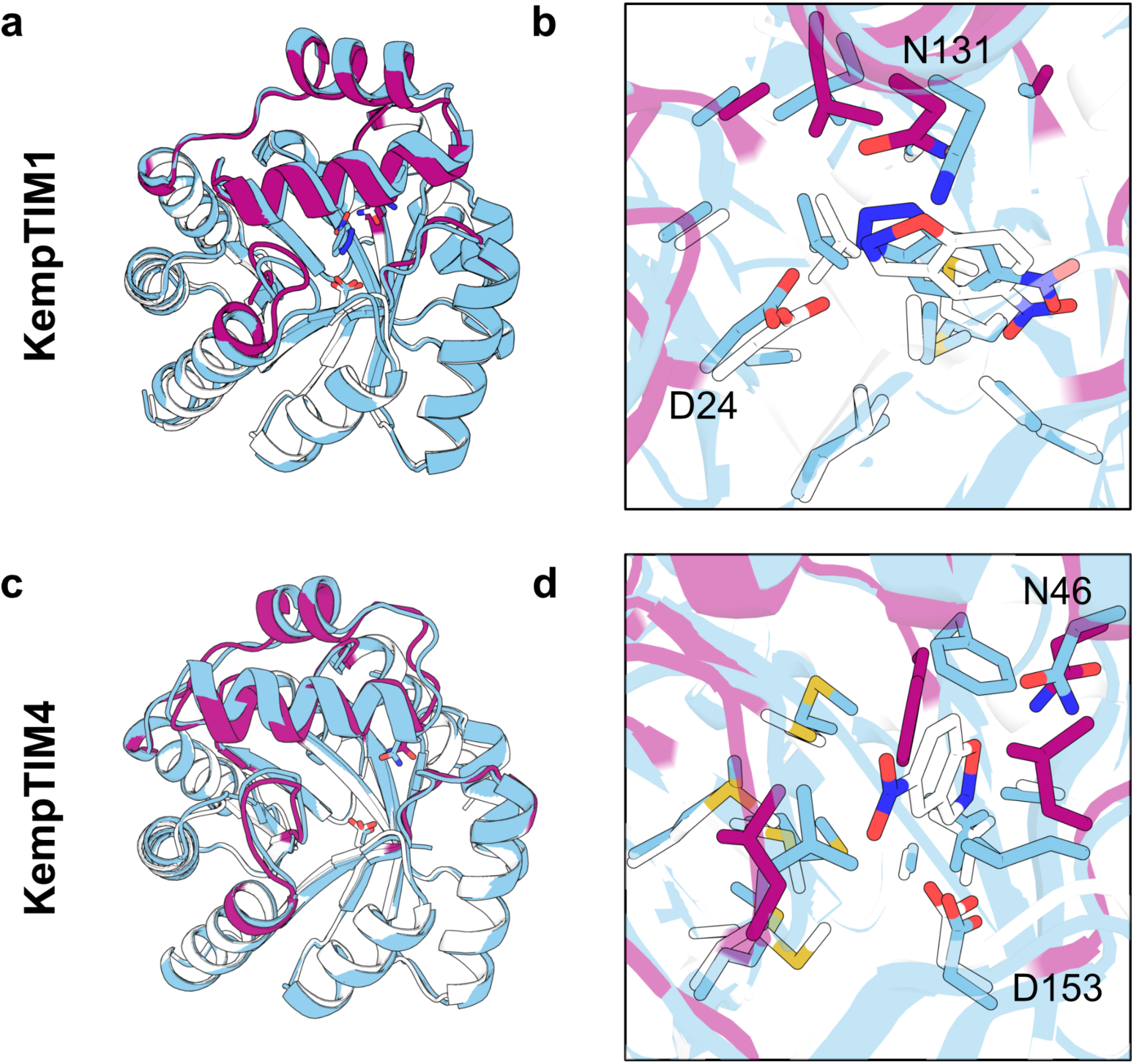
Crystal structures. Experimental structures are shown in blue and design models in white (minimal TIM barrel) and magenta (lid). (a) The crystal structure of KempTIM1 closely matches the design model, with backbone Cα root-mean-square deviations (RMSDs) of 0.68 Å for the full structure and 0.88 Å for the lid. (b) Active-site residues within the minimal TIM barrel of KempTIM1 more closely match their designed conformations than those in the lid, consistent with an ∼1 Å displacement of the lid relative to the TIM-barrel core. The co-crystallized transition-state analogue 6NBT adopts a binding pose similar to that in the design model, enabling hydrogen bonding to both the catalytic base D24 and the H-bond donor N131, despite N131 adopting a different rotamer in the crystal structure. (c) The crystal structure of KempTIM4 also shows close agreement with the design model (backbone Cα RMSD = 0.86 Å for the full structure and 1.34 Å for the lid). (d) Active-site residues of KempTIM4 show substantial divergence from the design model, potentially reflecting the absence of a bound transition-state analogue. The catalytic base D153 adopts a similar rotamer in both structures, whereas the catalytic H-bond donor N46 adopts a rotamer incompatible with transition-state stabilization, likely caused by the ∼2 Å displacement of the lid relative to the TIM-barrel core.

Despite crystallizing both proteins in the presence of the transition-state analogue 6-nitrobenzotriazole (6NBT), we observed clear electron density for this ligand only in the active site of KempTIM1 (Supplementary Figure 12), indicating weak binding in KempTIM4 under the crystallization conditions. In the KempTIM1 co-crystal structure (Figure 4b), active-site residues adopt conformations within 0.8 Å RMSD of the design model. The designed catalytic base D24 adopts a rotamer similar to that in the design model, enabling formation of a hydrogen bond with the bound transition-state analogue despite the ligand being slightly rotated relative to its designed orientation. This productive conformation of D24 is stabilized by an ordered water molecule (Supplementary Figure 12, Supplementary Figure 13). By contrast, the H-bond donor N131 adopts an alternative rotamer yet still forms a hydrogen bond with 6NBT, consistent with the observed sixfold decrease in catalytic efficiency when this residue is mutated to alanine (Figure 3c, Table 1). Notably, in the apo structure of KempTIM1 (Supplementary Figure 14), N131 adopts a rotamer that more closely matches the designed conformation and exhibits a Cα displacement of 1.1 Å relative to the co-crystal structure, placing it near its location in the design model. This backbone shift indicates a lid rearrangement upon substrate binding, a behavior commonly observed in natural enzymes. This conformational adjustment is accompanied by a rotameric change in the neighbouring residue R130, whose side chain reorients toward the binding pocket in the co-crystal structure and forms a water-mediated hydrogen bond with the transition-state analogue (Supplementary Figure 13).

The KempTIM1 co-crystal structure further shows a deviation of the transition-state analogue nitro group. Notably, the catalytic residues form hydrogen bonds that better accommodate a tautomeric form of 6NBT (Supplementary Figure 12, Supplementary Figure 15) corresponding to the 6-nitrobenzisoxazole substrate rather than the 5-nitro regioisomer targeted in the design. To determine whether this binding preference translates into altered regioselectivity, we measured kinetic parameters using 6-nitrobenzisoxazole as substrate. KempTIM1 exhibits an approximately twofold higher *k*_cat_/*K*_M_ for 6-nitrobenzisoxazole compared to the 5-nitro isomer (Supplementary Figure 16). These findings are consistent with the relatively large active-site pocket of KempTIM1, which would accommodate both regioisomers.

In contrast to KempTIM1, most binding pocket residues in KempTIM4 adopt conformations that differ from those intended to stabilize the transition state, indicating limited preorganization of the active site (Figure 4d). While the catalytic base D153 adopts the designed rotamer, the H-bond donor N46 adopts a conformation incompatible with transition-state stabilization. Consistent with this structural observation, mutation of N46 to alanine results in only a minor decrease in catalytic activity (Figure 3c, Table 1). Together, these structural features likely contribute to the orders-of-magnitude lower catalytic efficiency of KempTIM4 relative to KempTIM1 and provide clear targets for further optimization.

## Catalytic efficiency improved by ensemble-based design

To correct active-site deficiencies observed in KempTIM4, we applied an ensemble-based design^23,29^ strategy (Supplementary Figure 17). This approach uses protein backbones generated by crystallographic ensemble refinement^30^ as design templates. It can reveal conformational substates that better accommodate the transition state and its catalytic interactions, thereby enabling the accurate construction of enhanced active sites^23^.

During ensemble-based design, the identity of the catalytic base was fixed as Asp but its position within the β-barrel was allowed to vary (Methods). The identity of the H-bond donor was permitted to vary between Asn and Gln but could only be placed within the designed lid. Following sequence design with Triad, we filtered the design models (Methods) and predicted 6NBT-bound structures using the generative AI model Boltz-2^31^ (Supplementary Figure 18). This filtering pipeline yielded eight redesigned variants (KempTIM4a–h) comprising one of three distinct catalytic motifs (Supplementary Table 1, Supplementary Table 4, Supplementary Figure 4), none of which reproduced the motif found in the parent KempTIM4 (Table 1).

All second-round designs expressed solubly (Supplementary Table 2) and exhibited increased catalytic efficiency relative to KempTIM4 at pH 7, with two variants displaying improvements of at least two orders of magnitude (Supplementary Table 4, Supplementary Figure 8). The most active design, KempTIM4b, expressed as a stable monomeric TIM barrel (Figure 2) and reached a *k*_cat_/*K*_M_ of 3100 M^−1^ s^−1^ (Figure 3a), which is a >400-fold improvement thereby surpassing the most active first-round design KempTIM1 (Table 1). Mutation of the catalytic base of KempTIM4b abolished activity (Figure 3b), whereas substitution of the designed H-bond donor with alanine reduced catalytic efficiency approximately fivefold (Figure 3c). This result contrasts sharply with the parent KempTIM4, in which mutation of the H-bond donor had little effect on activity. Neither mutation altered the overall protein fold (Supplementary Figure 9). Like KempTIM1, KempTIM4b also showed enhanced activity at alkaline pH, with an apparent p*K*_a_ of 9.0 (Supplementary Figure 10, Supplementary Table 5). At pH 10, *k*_cat_ increased to 26 s^−1^ and *k*_cat_/*K*_M_ improved >1600-fold reaching 32,000 M^−1^ s^−1^ (Table 1, Figure 3d).

Notably, KempTIM4b reacted more efficiently with the intended target substrate 5-nitrobenzisoxazole than with 6-nitrobenzisoxazole, exhibiting 5- or 8-fold higher *k*_cat_/*K*_M_ at pH 7 and 10, respectively (Supplementary Table 4, Supplementary Figure 16). Together, the high catalytic efficiency and substrate selectivity of KempTIM4b demonstrate that *de novo* TIM barrels equipped with CANVAS-designed lids can host fully functional, selective active sites, and provide a platform for further optimization toward even more powerful catalysts.

## Active-site dynamics distinguish high- and low-activity designs

To probe the origins of activity differences among the designed enzymes, we performed molecular dynamics simulations of all active KempTIMs bound to the transition-state analogue 6NBT, using their design models as starting structures. Across all designs, the lids were more flexible than the minimal TIM-barrel core, but the helix anchoring the designed H-bond donor remained comparatively rigid (Supplementary Figure 19). Notably, this helix was most rigid in the highly active first-round design, KempTIM1, suggesting that increased helical rigidity may help stabilize the substrate and maintain catalytically productive geometry.

To test this hypothesis, we evaluated the ability of designed active sites to stabilize the transition-state analogue in a catalytically competent pose. For each trajectory snapshot, we calculated both the distance and angle of the two designed catalytic contacts (Figure 5a). The most active variants displayed narrow distributions centred on ideal values^32^ (Supplementary Figure 20), whereas lower-activity variants showed broadened distributions for one or both contacts. The H-bond donor interaction was generally the least consistently formed, in agreement with the observed lid displacement in the crystal structures (Figure 4) as well as the mutational studies (Figure 3b,c).

**Figure 5.**
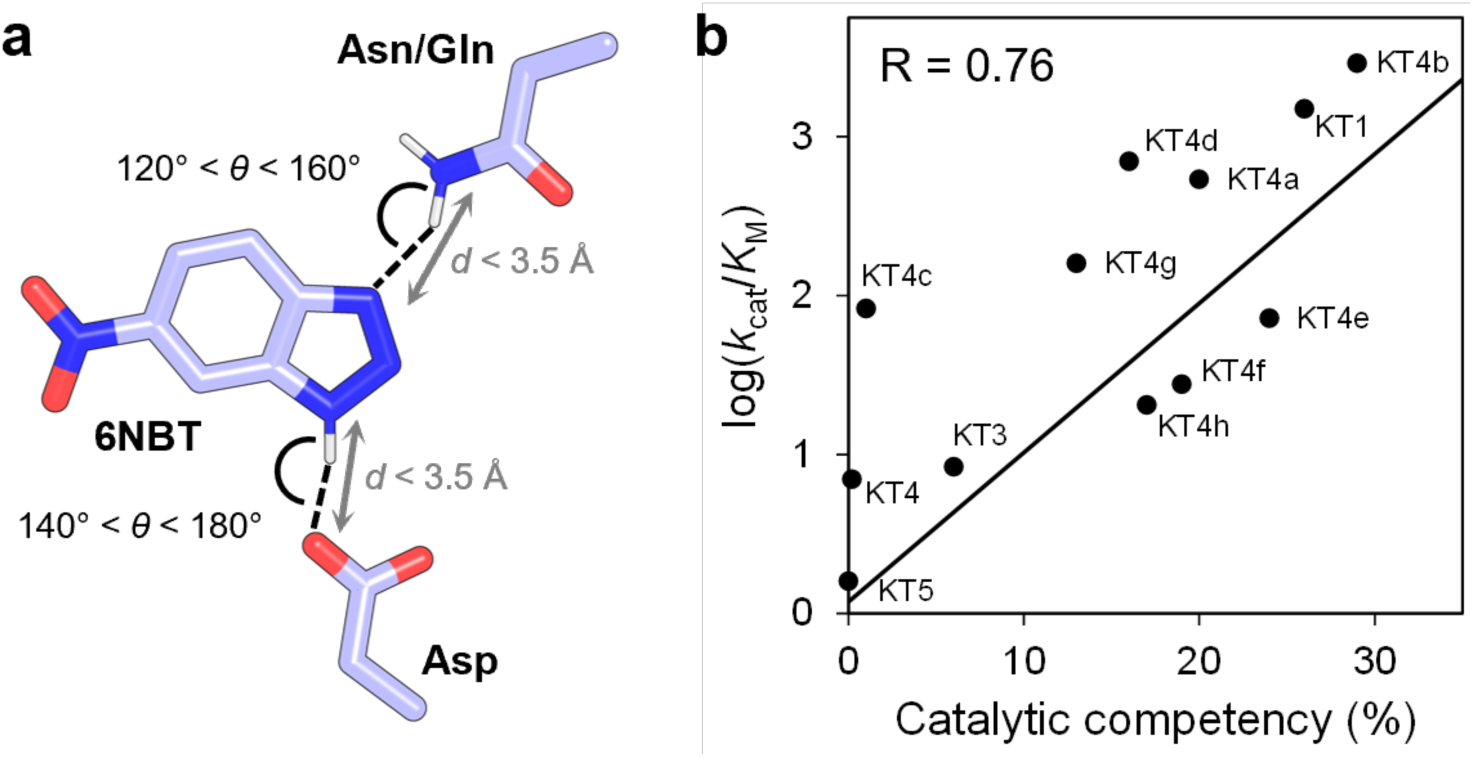
MD-derived catalytic competency correlates with experimental activity. (a) Geometric definitions of the catalytic contacts between the transition-state analogue (6NBT) and the designed general base (Asp) and H-bond donor (Asn/Gln). Catalytic angle (*θ*) and heavy-atom distance (*d*) thresholds defining competent contacts are indicated. (b) Correlation between catalytic-competency score, defined as the fraction of MD snapshots in which both catalytic contacts simultaneously satisfy the geometric criteria, and experimental catalytic efficiencies (*k*_cat_/*K*_M_). KempTIM (KT) variants with higher catalytic competency display higher activities (R = 0.76), with KempTIM1 and KempTIM4b ranking highest.

To assess whether differences in the ability of active sites to stabilize the designed catalytic contacts were related to the experimentally measured catalytic efficiencies, we calculated a catalytic competency score^23^, defined as the fraction of simulation snapshots in which both catalytic contacts satisfied geometric criteria expected for efficient catalysis. Catalytic competency scores correlated with measured *k*_cat_/*K*_M_ values across the variant set (R = 0.76, Figure 5b), with the two most active enzymes, KempTIM1 and KempTIM4b, exhibiting the highest scores. These findings indicate that this metric may serve as a useful in-silico filter for prioritizing future enzyme designs prior to experimental testing.

## Opportunities for CANVAS design

Natural TIM barrels use lids to bind substrates and form active sites conducive to catalysis^33^. By contrast, minimal *de novo* TIM barrels lack such lids and are inherently functionless. Using our CANVAS approach, we demonstrate that custom lid construction can transform these inert scaffolds into efficient enzymes. By designing various lids to simultaneously anchor a catalytic residue and form a substrate binding pocket, we created a series of *de novo* Kemp eliminases. One of these, KempTIM1, ranks among the most active *de novo* Kemp eliminases whose active site was obtained from a single round of computational design. Notably, its catalytic efficiency is 7-fold higher than that of the most active Kemp eliminase generated using scaffolds assembled combinatorially from fragments of homologous TIM-barrel templates^27^. This result demonstrates that minimal *de novo* TIM barrels equipped with custom-designed lids can yield enzymes that outperform those generated using fragments derived from natural scaffolds. Even variants with modest activity, such as KempTIM4, perform on par with many first-generation enzymes produced by conventional theozyme-based methods^4,6,34^, despite the added complexity of engineering a tailored lid comprising multiple loops to create an active site cavity. Importantly, CANVAS designs are evolvable. Using the crystal structure of KempTIM4 for ensemble-based design^23^, we increased catalytic efficiency by more than two orders of magnitude. Further improvements to activity of KempTIMs might be achieved using experimental optimization, for example NMR-guided directed evolution^35^, which has previously uncovered distal mutations that boost Kemp elimination rates in *de novo* enzymes to values approaching the diffusion limit^28^.

CANVAS also addresses a fundamental limitation of *de novo* enzyme design strategies. Unlike traditional approaches that retrofit catalytic groups into pre-existing folds^4,6^, CANVAS enables the modular customization of lids to satisfy the geometric and chemical requirements of virtually any reaction. This design versatility becomes more critical as the number of catalytic residues grows, a regime where fixed scaffolds make optimization increasingly difficult. While recent generative-AI methods address this problem by building entirely new folds around catalytic motifs extracted from the PDB or theozymes^36,37^, yielding functional *de novo* enzymes, the resulting proteins often bear little resemblance to natural enzymes. CANVAS offers a complementary strategy: it retains a familiar, well-behaved scaffold while precisely tailoring only the features essential for catalysis.

In the context of TIM barrels, this approach maintains a fold that is inherently versatile and widely used in nature across diverse chemistries. The β-barrel provides a stable framework for anchoring one or more catalytic residues, whereas the lid offers a tuneable element for controlling substrate binding and anchoring additional catalytic residues. Previous enzyme-design efforts have sought to harness this versatility by repurposing natural TIM barrels^4–6^ or by recombining natural barrel fragments *in silico*^27^, approaches that inherently restrict design space to existing lid architectures. By contrast, CANVAS decouples lid design from natural precedent, enabling the creation of bespoke lids tailored to specific catalytic demands. As a result, CANVAS transforms minimal, and even purely computationally generated, TIM barrels into genuine blank canvases for enzyme construction. We anticipate that this approach will unlock the full catalytic potential of designed TIM-barrel proteins across diverse chemical transformations. More broadly, the principles underlying CANVAS should extend beyond TIM barrels to other minimal protein architectures^38,39^, where designed lids could be used to configure and stabilize one or more catalytic residues for new-to-nature catalysis.

## Methods

### Reagents and solutions

All experiments were conducted using analytical-grade chemicals, with solutions prepared using double-distilled water.

### Circular permutation of DeNovoTIM6-SB with inpainting

Circular permutation of DeNovoTIM6-SB (PDB ID: 7OSV) was performed using the inpainting method described by Wang et al.^25^ The first residue of either the first or third β-strand was selected as the new N-terminus, with the corresponding βα-loop deleted (Supplementary Figure 2). A new connection for the corresponding βα-loop was modeled by inpainting 5 to 10 residues. Unresolved residues in the crystal structure were remodeled, keeping intact the positions and identities of the remaining residues. For each new connection, 100 designs were modeled, followed by relaxation and scoring using the Rosetta protein design software^40^. Structures of each design were predicted using AlphaFold2^22^ with model 4 weights. Designs with a pLDDT value above 90 and a Rosetta score below 3 Rosetta energy units per residue were kept. One design from each connection was selected for experimental characterization (Supplementary Table 1). The proteins were named NT6-CP1 (N-terminus on the first β-strand) and NT6-CP2 (N-terminus on the third β-strand).

### Computational enzyme design

All calculations were performed with the Triad protein design software^18^ (Protabit, Pasadena, CA, USA). Rotamer optimization was performed using a Monte Carlo with simulated annealing search algorithm. Input structures were prepared for Triad calculations via the *addH.py* application within Triad. The Kemp elimination transition state (TS) structure was built using parameters from Privett *et al.*^6^. To provide sidechain conformations, a backbone-independent conformer library (bda-bbind_1.0)^19^ was used for theozyme placement, and a backbone-independent rotamer library (bbind02.May.e2)^41^ with expansions of ± 1 standard deviation around χ_1_ and χ_2_ was used for active-site repacking. Energies were calculated using a modified version of the Phoenix energy function^18,42^ consisting of a Lennard-Jones 12–6 van der Waals term from the Dreiding II force field^43^ with atomic radii scaled by 0.9, a direction-dependent hydrogen bond term with a well depth of 8.0 kcal mol^−1^ and an equilibrium donor–acceptor distance of 2.8 Å,^44^ and an electrostatic term modeled using Coulomb’s law with a distance-dependent dielectric of 10. An energy benefit of –100 kcal mol^−1^ was applied when TS-side-chain interactions satisfied catalytic contact geometries (Supplementary Table 7), as described by Lassila *et al.*^19^

### Theozyme placement

AlphaFold2 predictions of NT6-CP1 and NT6-CP2, along with the 7MCD crystal structure, were used as backbone templates for theozyme placement. Amino-acid residues with a C_α_–C_β_ vector oriented towards the β-barrel interior and located at the C-terminus of each β-strand were selected as positions for introduction of the catalytic base (Asp). Neighbouring residues were mutated to alanine to avoid steric clashes with the TS (Supplementary Table 8). TS poses were built using the contact geometries listed in Supplementary Table 9, and those where the isoxazolic oxygen pointed toward the catalytic face of the TIM barrel and remained coplanar with the catalytic base were selected (Supplementary Figure 1). In parallel, the catalytic H-bond donor (Asn or Gln) was placed on a Gly-Asn/Gln-Gly tripeptide scaffold, and TS poses were built using H-bond donor side-chain contacts specified in Supplementary Table 9. The resulting TS was superimposed with the TS from the catalytic base calculation to create the full theozyme, positioning the catalytic base within the minimal TIM barrel and the H-bond donor in the empty space above the catalytic face. The Gly residues from the tripeptide were then deleted, preserving only the Asn/Gln residue, whose alpha carbon atom served as an anchor for the designed lid.

### Lid design

For all theozyme placements, peptide connections between the catalytic H-bond donor residue and the minimal TIM barrel structures were designed using RFdiffusion^20^. For each position of the H-bond donor alpha carbon, various βα-loops were selected as insertion points for a primary peptide fragment, and multiple fragment lengths were sampled to connect to the catalytic residue while preserving the TIM-barrel topology. Additional βα-loops were sometimes elongated to improve interactions with the primary fragment (Supplementary Figure 3). For each inserted primary fragment, 100–200 structures were sampled. Primary fragments lacking secondary structure, causing steric clashes with the TS, or displacing the H-bond donor from its theozyme position, were discarded (Supplementary Figure 1). The filtered fragments were sequence-optimized using ProteinMPNN^21^ using a temperature factor of 0.1, ensuring the minimal TIM barrel template sequence remained unchanged. Cysteine and methionine residues were excluded during sequence design. Structures of designed sequences were predicted using AlphaFold2^22^ or ColabFold v1.3.0^45^ with all five model weights, and those displaying an average pLDDT value > 90 and backbone RMSD < 3 Å relative to the RFdiffusion model were selected for active-site repacking.

### Active-site repacking

AlphaFold2 models generated above were used for a theozyme placement step^19^, as described previously, to verify that contacts between catalytic residues and the TS could be formed on these scaffolds. Active-site repacking calculations were then performed on the structures with this new theozyme placement. In these calculations, the TS was translated by ± 0.4 Å along each Cartesian coordinate in 0.2-Å steps and rotated about all three axes (origin at the TS geometric center) in 5° increments over a 10° range (clockwise and counterclockwise). This resulted in a total combinatorial search space of 15,625 possible poses. Residues near the catalytic amino acids and TS were designated as design positions. During repacking, these positions were allowed to sample rotamers of various amino acids, favoring hydrophobic and aromatic residues, while the identities of the catalytic residues remained fixed (Supplementary Table 10).

### Computational library design

After repacking, the CLEARSS computational library design algorithm^46^ was used to generate a combinatorial library comprising the most favorable amino acids predicted by Triad at each designed active-site position. In this method, libraries of a specific size configuration are generated from a pre-scored list of sequences using the highest probability set of amino acids at each position based on the sum of their Boltzmann weights. Using the list of energy-ranked sequences from active-site repacking as input, libraries of 192 sequences were generated. Rotamer configurations for each sequence in the library were optimized using the *cleanSequences.py* application within Triad to find the lowest-energy conformation of each sequence on its respective backbone, generating “cleaned” structures. To compare energies, the energy difference between each “cleaned” structure and a corresponding all-Gly structure in the absence of the TS was calculated. These energies are reported in Supplementary Table 11. Structures were cleaned with and without the TS to evaluate preorganization, as described previously^23^.

### Design filtering

Energies of the TS-bound and unbound “cleaned” structures were calculated using Triad. Dihedral angles of catalytic residues relative to the TS, as well as the solvent-accessible surface area (SASA) of the TS, were analyzed using PyMOL (v2.3.0, Schrödinger, LLC). Bottleneck radii of active-site entrances were evaluated using CAVER v3.0^47^, and the number of residues mutated to glycine or methionine was counted. Designs were filtered based on the following criteria (Supplementary Table 11): energy difference between bound and unbound structures below 0 kcal mol^−1^, catalytic residue dihedral angles outside 50° to 130° and –50° to –130°, SASA of TS between 80 and 200 Å², < 6 Met and < 4 Gly, bottleneck radius of active-site entrance > 0.9 Å, and < 7 non-preorganized residues. Designs passing all criteria were used for structure prediction with AlphaFold2 or ColabFold (v1.3.0), where designs with a backbone RMSD < 3 Å relative to the structure cleaned by Triad, overall pLDDT > 90, and lid pLDDT > 85 were chosen for experimental characterization (Supplementary Table 1).

### Sequence optimization of variants 1–5 for enhanced expression

Sequence optimization of variants 1–5 was performed using ProteinMPNN with LigandMPNN weights^48^. Amino-acid identities of all residues designed during active-site repacking with Triad were retained. Four sequences were generated using the AlphaFold2 models of each variant as a template, omitting cysteine. The generated sequences were used for structure prediction with ColabFold^45^ using all five model weights. Predictions were filtered based on their all-atom RMSD to the input AlphaFold2 model. Theozyme placement was carried out using ColabFold structures, as previously described, to ensure that catalytic contacts were preserved. A single design for each variant was selected for experimental characterization (named KempTIM1–5, Supplementary Table 1).

### Ensemble-based design

Two ensembles of backbone templates were generated from the KempTIM4 crystal structure (PDB ID: 9QKX) using the ensemble refinement protocol^30^ implemented in the PHENIX macromolecular structure determination software. In ensemble refinement, local atomic fluctuations are sampled using molecular dynamics simulations accelerated and restrained by electron density to produce ensemble models fitted to diffraction data. Briefly, input crystal structures were edited to remove low-occupancy conformers and assign an occupancy of 1.0 to the remaining conformer. Following addition of riding hydrogens, parallel ensemble refinement simulations were performed using various combinations of the parameters p_TLS_ (0.6, 0.8, 0.9, 1.0), τ_x_ (0.5, 1.5, 2.0) and w_x-ray_ (2.5, 5.0, 10.0), where p_TLS_ describes the percentage of atoms included in a translation-libration-screw (TLS) model use to remove the effects of global disorder, τ_x_ is the simulation time-step and w_x-ray_ is the coupled tbath offset, which controls the extent to which the electron density contributes to the simulation force field such that the simulation runs at a target temperature of 300 K. Two ensembles were combined to form a 60-member ensemble (Supplementary Table 12).

Theozyme placement was performed as described previously on each ensemble member, targeting positions for the base and H-bond donor indicated in Supplementary Table 13. LigandMPNN was used to predict active-site mutations, allowing only hydrophobic and aromatic residues. Amino acids with a mean probability >1% across ten batches were selected as the sequence space for active-site repacking in Triad, performed as described above. Triad models (Supplementary Figure 4) and Boltz-2 predictions (Supplementary Figure 18) were used to filter the designs (Supplementary Table 14 and Supplementary Table 15). Designs were retained only if Boltz-2 models reproduced the intended catalytic contacts and the predicted ligand binding pose deviated by < 1.0 Å root-mean-square deviation from the corresponding Triad model.

### Cloning and generation of constructs

Genes for all proteins were ordered as codon-optimized fragments flanked by restriction sites for *Nde*I and *Xho*I from Twist Bioscience (South San Francisco, CA, USA). After digestion with *Nde*I and *Xho*I, the gene fragment was ligated into pET21b(+) for variants 1–5 and KempTIM1–5, or pET29b(+) for KempTIM6–9. Catalytic knockout mutations were introduced by a modified QuikChange PCR protocol utilizing KAPA Polymerase followed by *Dpn*I digestion. *E. coli* Top10 or BL21 (DE3) cells were transformed with ligated vector or reaction mixture and plated on Lysogeny Broth (LB) agar plates containing 100 µg mL^−1^ ampicillin or 50 µg mL^−1^ kanamycin as selection markers. Single colonies were used to inoculate 5 mL LB supplemented with 100 µg mL^−1^ ampicillin or 50 µg mL^−1^ kanamycin. After overnight growth (37 °C, 250 rpm), cells were harvested by centrifugation and DNA was isolated using NucleoSpin Plasmid EasyPure-Kit (Machery & Nagel) according to the manufacturer’s protocol. Vector assembly and introduction of mutations were validated by sequencing (Eurofins Genomics) using standard T7 primers.

### Protein expression and purification

*E.coli* BL21 (DE3) cells (Novagen) were transformed with plasmid, plated on agar plates containing antibiotic, and incubated overnight at 37 °C. Single colonies were picked to inoculate precultures and incubated at 30–37 °C overnight. On the next day, 1 L LB supplemented with antibiotic was inoculated with 5–10 mL of preculture and incubated at 37 °C until OD600 reached a value between 0.6 and 0.8. Overexpression was induced by adding isopropyl-β-thiogalactoside (IPTG) to a final concentration of 0.1 mM for variants 1–5 and KempTIM1–5 or 1 mM for KempTIM6–9 followed by overnight incubation at 16–20 °C with shaking. Cells were harvested by centrifugation and pellets were either frozen at –20 °C or used directly for purification.

Two different purification schemes were employed. In the first scheme, cell pellets were resuspended in 35 mL buffer A (50 mM sodium phosphate pH 7.0, 100 mM NaCl, 10 mM imidazole). Resuspended cells were lysed by sonication (Branson Ultrasonic Sonifier 250, output 4, duty cycle 40%, 3 × 3 min) and centrifuged. The supernatant was loaded onto a HisTrapHP column (5 mL, Cytiva Life Science) equilibrated with buffer A and coupled to an ÄKTApure system (Cytiva Life Science). After washing with 10 column volumes (CV) of buffer A, the protein was eluted with a linear gradient over 20 CV to 60 % buffer B (50 mM sodium phosphate pH 7.0, 100 mM NaCl, 500 mM imidazole). For crystallisation of KempTIM4-gsg, which contains an additional GSG tripeptide linker between its His-tag and first TIM barrel residue (Supplementary Table 1) and KempTIM1, an additional overnight TEV-cleavage step was performed during a dialysis against buffer C (50 mM of sodium phosphate pH 7.0, 100 mM NaCl). The dialyzed protein was loaded onto a HisTrapHP column (5 mL, Cytiva Life Science) equilibrated with buffer C and coupled to an ÄKTApure system (Cytiva Life Science) and the flowthrough was collected. Fractions containing the protein were pooled, concentrated with a centrifugal concentrator, and loaded onto a HiLoad 26/600 Superdex 75 preparative grade column (Cytiva Life Sciences) preequilibrated in buffer C and eluted with 1 CV buffer C.

In the second scheme, cell pellets were resuspended in 8 mL buffer D (5 mM imidazole in 100 mM potassium phosphate buffer at pH 8.0) supplemented with 1 mg mL^−1^ lyophilized lysozyme (MP Biomedicals) and 1 U mL^−1^ benzonase nuclease (Merck Millipore). Cells were homogenized using an Avestin EmulsiFlex-B15 cell disruptor and centrifuged. Due to their high thermostability, greater purity for KempTIM4a-h could be achieved through heating the lysate at 70 °C for 1 hour prior to centrifugation. Proteins were purified from the lysate by immobilized metal affinity chromatography using Ni-NTA agarose (Qiagen) pre-equilibrated with buffer D in individual Econo-Pac gravity-flow columns (Bio-Rad). Contaminants were washed away using buffer E (10 mM imidazole in 100 mM potassium phosphate buffer pH 8.0) and then 20 mM imidazole in the same buffer. Proteins were eluted with 5 mL of buffer F (250 mM imidazole in 100 mM potassium phosphate buffer pH 8.0). Proteins were further subjected to gel filtration in buffer C using an ENrich SEC 70 size-exclusion chromatography column (Bio-Rad). Purity was confirmed by sodium dodecyl sulfate-polyacrylamide gel electrophoresis (SDS-PAGE) and fractions containing the protein of interest were pooled. Protein concentration was determined spectrophotometrically using the absorption at 280 nm and applying Beer-Lambert’s law using calculated extinction coefficients obtained from the ExPAsy ProtParam tool (https://web.expasy.org/protparam/).

### Far-UV circular dichroism (CD) spectroscopy

CD spectra were collected with a Jasco J-710 or J-815 spectrometer. Spectra were recorded using a 1-mm quartz cuvette at 20 °C with scanning speed 10–100 nm min^−1^, bandwidth 1 nm, response time 1 s. Experiments were performed in 20 mM sodium phosphate buffer (pH 7.0), or 50 mM sodium phosphate buffer (pH 8.0) supplemented with 100 mM sodium chloride, using a protein concentration of approximately 0.2 mg mL^−1^. For each protein, 3 to 10 spectra were accumulated and averaged. For data normalization, a buffer spectrum was subtracted and the signal was converted to mean residue molar ellipticity using [*θ*_MRE_] = *θ*/(n × *d* × *c*), where *n* is the number of residues in the protein, *θ* the collected ellipticity in mdeg, *d* the path length in mm, and *c* the protein concentration in M. Thermal denaturation assays were performed by heating the samples from 20 to 95 °C at a rate of 1 °C per minute and ellipticity at 222 nm was measured every 1 °C. Melting temperature (T_m_) was determined following the protocol by Greenfield^49^.

### Size-exclusion chromatography-multi angle light scattering

SEC-MALS measurements were performed as described in Beck *et al*.^12^ For all measurements, buffer C, an injection volume of 50 µL, and a protein concentration of 2 mg mL^−1^ were used.

### Steady-state kinetics

Assays were performed in sodium phosphate (pH 7.0), Tris (pH 7.0–8.5), or CHES (pH 9.0–10.0) buffers, all at 50 mM with 100 mM NaCl and 10% MeOH in a Spark (Tecan) or Synergy H1 (Biotek) plate reader. Enzyme concentrations varied from 0.2 to 80 µM. Reactions (200 μL final volume) were initiated by addition of varying concentrations (0.015–2 mM) of 5-nitrobenzisoxazole (abcr or AAblocks) or 6-nitrobenzisoxazole (AAblocks) dissolved in methanol (final methanol concentration 10%). Product formation was monitored at 380 nm (ε = 15,800 M^−1^ cm^−1^)^6^ or 400 nm (ε = 2,870 M^−1^ cm^−1^)^50^ for 5-nitrobenzisoxazole and 6-nitrobenzisoxazole, respectively, in individual wells of a 96-well plate (Nunc or Greiner Bio-One). Path lengths for each well were calculated ratiometrically using the difference in absorbance at 900 and 975 nm. Control measurements without protein were conducted for each substrate concentration and subtracted from the enzyme measurements. Linear phases of the kinetic traces were used to measure initial reaction rates. If the enzyme showed saturation within the substrate concentration used, the data were fitted to the Michealis-Menten model using (v_0_ = (v_max_ × [S]) / (*K*_M_ + [S]) where v_0_ is the initial velocity rate, v_max_ is the maximal velocity rate, [S] is the substrate concentration, *K*_M_ is the Michaelis constant. The catalytic constant *k*_cat_ was calculated with *k*_cat_ = v_max_ / [E] where [E] is the enzyme concentration. If saturation was not achieved within the substrate’s solubility limit, a linear equation (v_0_ = *k*_cat_/*K*_M_ × [S]) was used.

### Crystallization and structure determination

KempTIM4-gsg and KempTIM1 (Supplementary Table 1) were purified using the first purification scheme, with the final size-exclusion chromatography step performed in 20 mM HEPES buffer pH 8.0 supplemented with 20 mM NaCl. Crystallization screens were set up using the sitting-drop vapor diffusion method with a Phoenix pipetting robot (Art Robbins Instruments) and commercially available sparse-matrix screens (NeXtal) in 96-well sitting-drop plates (3-drop Intelli-Plates, Art Robbins Instruments). Protein, reservoir solutions were mixed in ratios of 1:1, 2:1, and 1:2 and crystals were grown at 293 K. The protein concentration of KempTIM4-gsg was 12 mg mL^−1^, supplemented with 5 mM 6NBT (5% DMSO), and diffracting crystals were obtained after 40 days in 4 M sodium formate. Cryoprotection was achieved by adding glycerol to a final concentration of 25%.

The protein concentration of KempTIM1 was 8.75 mg mL^−1^. For the inhibitor-bound complex structure 6NBT was added as a solid in excess and the solution was incubated for 1 h and subsequently centrifuged. Apo KempTIM1 crystals were obtained after 48 h in droplets containing 0.2 M ammonium acetate, 0.1 M Bis-Tris pH 5.5, 25% w/v PEG 3350 as reservoir solution. The 6NBT complex crystals were observed after 24 h in droplets with 0.1 M sodium acetate pH 4.5, 0.1 M Bis-Tris pH 5.5, 25% w/v PEG 3350 as reservoir. The KempTIM1 crystals were cryo-protected by adding 20% ethylene glycol to the corresponding reservoir solution. All crystals were mounted using cryo-loops on SPINE standard bases and flash-cooled in liquid nitrogen.

Diffraction data were collected at 100 K at beamline BL 14.1 with a Pilatus3 S 6M detector at the BESSY II synchrotron (Helmholtz-Zentrum Berlin). Data processing was performed using X-ray Detector Software APP3 (XDSAPP3)^51^ with XDS^52^, and data quality was assessed using phenix.xtriage^53^. Phases were solved by molecular replacement using AlphaFold2 predictions as search models with Phaser^54^. The resulting models were manually rebuilt using Coot^55^ and refined with phenix.refine^53^ in an iterative process.

The structure of KempTIM4-gsg was refined to a resolution of 2.3 Å. Despite crystallization with 6NBT, no electron density corresponding to the transition-state analogue was observed in the active site; instead, the density was best interpreted as a bound glycerol molecule from the cryoprotectant. The structures of KempTIM1 apo and co-crystallized with 6NBT were refined to resolutions of 1.25 Å and 1.20 Å, respectively. The high resolution of the co-crystal structure and the well-defined electron density of 6NBT in the active site enabled conclusive placement of the transition-state analogue. Additionally, there is also a diffuse electron density, which cannot be interpreted unambiguously, indicating weak binding of solvent molecules. However, a putative ethylene glycerol molecule was placed into the density to account for the residual density, as it was used as cryoprotectant. Refinement statistics and crystallographic data are shown in Supplementary Table 6. Coordinates and structure factors were validated and deposited in the PDB under accession codes 9TZD, 9U01 and 9QKX.

### Size-exclusion chromatography small-angle X-ray scattering (SEC-SAXS)

SEC-SAXS measurements were conducted at the BioSAXS beamline BM29 of the ESRF in Grenoble, France. A Superdex 75 Increase 10/300 GL column (Cytiva Life Sciences) was used with a flow rate of 0.8 mL min^−1^ in buffer C. KempTIM4-gsg was measured at a concentration of 5 mg mL^−1^, with an injection volume of 100 µL. Data processing and analysis were performed using ATSAS 3.2.1^56,57^. For analysis, AlphaFold2 models of KempTIM4 and NT6-CP2, as well as an AlphaFlow^58^ model of KempTIM4 with a displaced lid obtained using the base molecular dynamics weights were used. For each of these models a theoretical scattering curve were generated and fitted to the experimental data using CRYSOL^59^.

### Molecular dynamics (MD) simulations

The Amber 2020 software (http://ambermd.org/) with the ff19SB protein force field^60^, gaff2 ligand force field^61^, and OPC water model^62^ was used for all simulations. A cutoff of 10 Å was applied to electrostatics modeled using the particle mesh Ewald method^63^. All MD trajectories were run with a time step of 2 fs. Models of each design generated using Triad as described above were prepared for MD. MD trajectories were run for each enzyme in the presence and absence of bound transition-state analogue 6-nitrobenzotriazole (6NBT). Parameters for 6NBT were generated using the Antechamber package^64^. Hydrogen atoms were added using Reduce^65^, and the prepared structures were solvated with OPC water in a truncated octahedral box with periodic boundary conditions where the distance between the protein surface and the box edges was set to 10 Å. The addions2 algorithm in Amber was used to place counterions to neutralize the system. The system was minimized (solvent, protein, then whole system) using the steepest descent method with positional restraints of 500 kcal mol^−1^ Å^−2^. The system was then heated to a temperature of 300 K over 240 ps while restraints were gradually removed, followed by equilibration under an NPT ensemble at 300 K and 1 bar for 10 ns and subsequent equilibration under an NVT ensemble at 300 K for another 10 ns. Constant temperature and pressure were achieved using the Langevin thermostat^66^ and Berendsen barostat^67^, respectively. Following equilibration, ten replicas of 100-ns production simulations were run. For analyzing catalytic contacts, 1000 snapshots separated by 0.1 ns each were extracted from the production trajectory. Catalytic interactions and RMSF values were extracted using CPPTRAJ and PYTRAJ, respectively^68^.

## Acknowledgments

J.B. and B.H. acknowledge support from the Elite Network of Bavaria and its study program “Biological Physics”. B.H. acknowledges support from European Union’s Horizon 2020 research and innovation program under grant agreement No 951375 (ArtMotor). R.A.C. acknowledges grants from the Natural Sciences and Engineering Research Council of Canada (RGPIN-2021-03484 and RGPAS-2021-00017) and the Canada Foundation for Innovation (26503). B.J.S. was supported by an NSERC CREATE scholarship. This research was enabled in part by support provided by Compute Ontario (www.computeontario.ca) and the Digital Research Alliance of Canada (alliancecan.ca). We acknowledge financial support and allocation of synchrotron beamtime by HZB at BESSY and at ESRF in Grenoble and thank the beamline staff for support. We thank Sabrina Wischt for technical support, Janosch Hennig for help with SAXS data analysis and the University of Bayreuth Centre of International Excellence “Alexander von Humboldt” for facilitating this collaboration through a short-term grant to R.A.C.

