## Supplementary Information for "Customizing the Structure of Minimal TIM Barrels to Craft Efficient *De Novo* Enzymes"

[†] These authors contributed equally.

*** Corresponding authors**

**This file includes:**

Supplementary Tables 1–15

Supplementary Figures 1–20

**Supplementary Table 1.** Amino-acid sequences

| Enzyme | # residues | MW [a] (kDa) | Sequence [b] |
|---|---|---|---|
| **NT6-CP1** | 188 | 22.1 | RIAYRSDDWRDLQEALKKGADILIVDATDKDEAWKQVEILRRLGAKEIAYRSDDWRDLQEALKKGADILIVDATDPEEAWKQVEILRRLGAKRIAYRSDDWRDLQEALKKGADILIVDATDKDEAWKQVEILRRLGAKEIAYRSDDWRDLQEALKKGADILIVDASDAERALKQVEILRRLEHHHHHH |
| **NT6-CP2** | 188 | 22.1 | EIAYRSDDWRDLQEALKKGADILIVDATDPEEAWKQVEILRRLGAKRIAYRSDDWRDLQEALKKGADILIVDATDKDEAWKQVEILRRLGAKEIAYRSDDWRDLQEALKKGADILIVDADDPEEAEKQVEILRRLGAKRIAYRSDDWRDLQEALKKGADILIVDATDKDEAWKQVEILRRLEHHHHHH |
| **7MCD** | 191 | 21.4 | DILIVNPDDFEKGVEEVKELKRHGAKIIAYISKSAEELKKAEKAGADILIVNPDDFEKGVEEVKELKRHGAKIIAYISKSAEELKKAEKAGADILIVNPDDFEKGVEEVKELKRHGAKIIAYISKSAEELKKAEKAGADILIVNPDDFEKGVEEVKELKRHGAKIIAYISKSAEELKKAEKAGLEHHHHHH |
| **Variant 1** | 225 | 26.3 | EIAYASDDWRDLQEALKKGADILD**VVNVNP**EEAWKQVEILRRLGAKRILYASLRWRDLQEALKKGADILA**VPLVDK**DEAWKQVEILRRLGAKEIAYGSDDWRDLQEALKKGADILGV**ASNTPRVRRLVLR**N**LETLYSREEAERIVEEKIKLNDPE**EAEKQVEILRRLGAKRIMYASDDWRDLQEALKKGADILLV**AADDESEESK**DEAWKQVEILRRLEHHHHHH |
| **Variant 2** | 225 | 26.3 | EIAYASDDWRDLQEALKKGADILD**VVNVNP**EEAWKQVEILRRLGAKRIAYASLRWRDLQEALKKGADILA**VPLVDK**DEAWKQVEILRRLGAKEIAYGSDDWRDLQEALKKGADILGV**ASNTPRVRRLVLR**N**LETLYSREEAERIVEEKIKLNDPE**EAEKQVEILRRLGAKRIMYASDDWRDLQEALKKGADILLV**VADDESEESK**DEAWKQVEILRRLEHHHHHH |
| **Variant 3** | 227 | 25.6 | RIAYA**SLASRW**RDLQEALKKGADILMV**ALARDAASVRALAEGAAG**N**GDPRSVAELLALLRPTDKD**EAWKQVEILRRLGAKEIAYMSDDWRDLQEALKKGADILMV**LASSALSPEE**AWKQVEILRRLGAKRIAYGSGDWRDLQEALKKGADILD**VLGAKGKD**EAWKQVEILRRLGAKEIAYASFDWRDLQEALKKGADILMV**FAVLDA**ERALKQVEILRRLEHHHHHH |
| **Variant 4** | 227 | 25.6 | RIAYM**SLASRW**RDLQEALKKGADILMV**ALARDAASVRALAEGAAG**N**GDPRSVAELLALLRPTDKD**EAWKQVEILRRLGAKEIAYMSDDWRDLQEALKKGADILMV**LASSALSPEE**AWKQVEILRRLGAKRIAYGSGDWRDLQEALKKGADILD**VLGAKGKD**EAWKQVEILRRLGAKEIAYASFDWRDLQEALKKGADILVV**FAVLDA**ERALKQVEILRRLEHHHHHH |
| **Variant 5** | 227 | 25.6 | RIAYA**SLASRW**RDLQEALKKGADILMV**ALARDAASVRALAEAAAG**N**GDPRSVAELLALLRPTDKD**EAWKQVEILRRLGAKEIAYMSDDWRDLQEALKKGADILMV**LASSALSPEE**AWKQVEILRRLGAKRIAYGSGDWRDLQEALKKGADILD**VLGAKGKD**EAWKQVEILRRLGAKEIAYASFDWRDLQEALKKGADILMV**FAVLDA**ERALKQVEILRRLEHHHHHH |
| **Variant 6** | 211 | 23.5 | DILGVAPDDFEKGVEEVKELKRHGAKIIGYMSKSAEELKKAEKAGADILLVAPDDFEKGVEEVKELKRHGAKIIAYMSKSAEELKKAEKAGADILMVYPDDFEKGVEEVKELKRHGAKIIAYLSKSAEELKKAEKAGADILDV**ANAE**Q**EKIASKFLGRKTKVKIEENDF**EKGVEEVKELKRHGAKIIAYGSKSAEELKKAEKAGWHHHHHH |
| **Variant 7** | 211 | 23.5 | DILGVAPDDFEKGVEEVKELKRHGAKIIGYMSKSAEELKKAEKAGADILLVAPDDFEKGVEEVKELKRHGAKIIAYMSKSAEELKKAEKAGADILMVFPDDFEKGVEEVKELKRHGAKIIAYLSKSAEELKKAEKAGADILDV**ANAE**Q**EKIASKFMGRKTKVKIEENDF**EKGVEEVKELKRHGAKIIAYGSKSAEELKKAEKAGWHHHHHH |
| **Variant 8** | 211 | 23.5 | DILGVAPDDFEKGVEEVKELKRHGAKIIGYMSKSAEELKKAEKAGADILMVAPDDFEKGVEEVKELKRHGAKIIAYMSKSAEELKKAEKAGADILAVFPDDFEKGVEEVKELKRHGAKIIAYLSKSAEELKKAEKAGADILDV**ANAE**Q**EKIASKLMGRKTKVKIEENDF**EKGVEEVKELKRHGAKIIAYGSKSAEELKKAEKAGWHHHHHH |
| **Variant 9** | 211 | 23.5 | DILGVAPDDFEKGVEEVKELKRHGAKIIGYMSKSAEELKKAEKAGADILLVAPDDFEKGVEEVKELKRHGAKIIAYMSKSAEELKKAEKAGADILMVYPDDFEKGVEEVKELKRHGAKIIAYLSKSAEELKKAEKAGADILDV**ANAE**Q**EKIASKLLGRKTKVKIEENDF**EKGVEEVKELKRHGAKIIAYGSKSAEELKKAEKAGWHHHHHH |
| **KempTIM1** | 235 | 25.2 | HHHHHHENLYFQSGSGEIAYASGDADHLLAAREAGADILD**VVDVDP**ARALAQVRRLRAAGARRILYASLRVDDLLAALEAGADILA**VPDLDH**DAALAQIRALKAAGAREIAYGSPDADHLLEAREAGADILGV**AADTPAVRELWLR**N**LRQVFSEEEAREILDRRIHLPDPA**TAIEQVRRLRAAGAKRIMFASDDVDHLAAAKRAGADILLV**AEAGGSAEAR**AAALAQVRRLKALW |
| **KempTIM2** | 235 | 25.6 | HHHHHHENLYFQSGSGEIAFASGDVDHLLAAMEAGADILD**VVNVDP**AAGLEQVRRLKAAGAKRIAYASLRVEDLLAALKAGADILA**VPDYDH**ERELRIIRELKRAGAKEIAYGSSDADHLLEARRAGADILGV**AADTPAVRELWLN**N**LRQVFSEEEARRILDERIHLPPPA**EAIEQVRRLRAAGAERIMYASDDVDHLAAAKEAGADILLV**VEAGGSAEAL**ERALAQVRRLKALW |
| **KempTIM3** | 238 | 25.3 | HHHHHHENLYFQSGSGRIAYA**ALASDL**DSLVEALKLGADILMV**ALMADAAAVRAMAEGLHA**N**GDPRSVAELEALLRPTDLD**DALAAVRELKALGAKEIAFMSHDVDHLIRAMEAGADILMV**LESSATSVEA**ALAQVRRLKAAGAKRISFGSGDVAHLKAAMEAGADILD**VLERHGLD**VALAQIRELKAAGAKEIAFASLDPDHLLRAREEGADILMV**FGATDP**ARALATVRYLRAQAW |

| | | | |
|---|---|---|---|
| **KempTIM4** | 233 | 25.0 | HHHHHHENLYFQSRIAYM**ALASDL**DSLVEALKLGADILMV**ALMADAAAVRAMAEGLHA**N**GDPRSVAE**<br>**LEALLRPTDLD**DALAAVRELKALGAKEIAFMSHDVDHLIRAMEAGADILMV**LESSATSVEA**ALAQVR<br>RLKAAGAKRISFGSGDVAHLKAAMEAGADILD**VLERHGLD**VALAQIRELKAAGAKEIAFASLDPDHL<br>LRAREEGADILVV**FGATDP**ARALATVRYLRAW |
| **KempTIM5** | 238 | 25.1 | HHHHHHENLYFQSGSGRIALA**ALAADE**DTLLRALELGADILMV**ALLPDAAAVRAAAIAAHA**N**GDPRS**<br>**VEELEALLRPTDVD**AALAVVRRLKAAGAKEIAFMSHDVDHLIAAMEAGADILMV**LASVATSVEA**ALE<br>QVRRLKAAGAKRIAFGSGDVDELIAAMEAGADILD**VLEARGLD**VALAQIRRLKEAGAKEIAFASFDA<br>AHLLEARRAGADILMV**FALLDP**DRALAIVEELRAAAW |
| **KempTIM4a** | 233 | 24.8 | HHHHHHENLYFQSRIAYI**ALASDL**DSLVEALKLGADILAV**LLMADAAAVRALAE**Q**LHAIGDPRSVAE**<br>**LEALLRPTDLD**DALAAVRELKALGAKEIAFASHDVDHLIRAMEAGADILAV**VESSATSVEA**ALAQVR<br>RLKAAGAKRILFGSGDVAHLKAAMEAGADILD**VLARHGLD**VALAQIRELKAAGAKEIAFVSLDPDHL<br>LRAREEGADILYV**VGATDP**ARALATVRYLRAW |
| **KempTIM4b** | 233 | 24.8 | HHHHHHENLYFQSRIAYI**ALASDL**DSLVEALKLGADILAV**LLMADAAAVRALAE**Q**LHAIGDPRSVAE**<br>**LEALLRPTDLD**DALAAVRELKALGAKEIAFASHDVDHLIRAMEAGADILAV**VESSATSVEA**ALAQVR<br>RLKAAGAKRILFASGDVAHLKAAMEAGADILD**VLARHGLD**VALAQIRELKAAGAKEIAFVSVDPDHL<br>LRAREEGADILFV**LGATDP**ARALATVRYLRAW |
| **KempTIM4c** | 233 | 25.0 | HHHHHHENLYFQSRIAYD**ALASDL**DSLVEALKLGADILLV**ALMADAAAVRVWAEM**Q**HLAGDPRSVAE**<br>**LEALLRPTDLD**DALAAVRELKALGAKEIAFLSHDVDHLIRAMEAGADILMV**VEVSATSVEA**ALAQVR<br>RLKAAGAKRIMFASGDVAHLKAAMEAGADILA**VAERHGLD**VALAQIRELKAAGAKEIAFASMDPDHL<br>LRAREEGADILIV**AGATDP**ARALATVRYLRAW |
| **KempTIM4d** | 233 | 25.1 | HHHHHHENLYFQSRIAYA**ALASDL**DSLVEALKLGADILIV**LLMADAAAVRAMAEMLHL**N**GDPRSVAE**<br>**LEALLRPTDLD**DALAAVRELKALGAKEIAFMSHDVDHLIRAMEAGADILMV**FELSATSVEA**ALAQVR<br>RLKAAGAKRIAFDSGDVAHLKAAMEAGADILM**VAERHGLD**VALAQIRELKAAGAKEIAFVSFDPDHL<br>LRAREEGADILVV**AGATDP**ARALATVRYLRAW |
| **KempTIM4e** | 233 | 25.0 | HHHHHHENLYFQSRIAYA**ALASDL**DSLVEALKLGADILIV**ILMADAAAVRAYAELLHL**N**GDPRSVAE**<br>**LEALLRPTDLD**DALAAVRELKALGAKEIAFLSHDVDHLIRAMEAGADILMV**LESSATSVEA**ALAQVR<br>RLKAAGAKRIAFDSGDVAHLKAAMEAGADILM**VAERHGLD**VALAQIRELKAAGAKEIAFASFDPDHL<br>LRAREEGADILVV**AGATDP**ARALATVRYLRAW |
| **KempTIM4f** | 233 | 25.0 | HHHHHHENLYFQSRIAYA**ALASDL**DSLVEALKLGADILLV**ILMADAAAVRAYAEMLHL**N**GDPRSVAE**<br>**LEALLRPTDLD**DALAAVRELKALGAKEIAFLSHDVDHLIRAMEAGADILLV**LESSATSVEA**ALAQVR<br>RLKAAGAKRISFDSGDVAHLKAAMEAGADILM**VAERHGLD**VALAQIRELKAAGAKEIAFASFDPDHL<br>LRAREEGADILVV**AGATDP**ARALATVRYLRAW |
| **KempTIM4g** | 233 | 24.9 | HHHHHHENLYFQSRIAYA**ALASDL**DSLVEALKLGADILIV**VLMADAAAVRAFAELLHL**N**GDPRSVAE**<br>**LEALLRPTDLD**DALAAVRELKALGAKEIAFLSHDVDHLIRAMEAGADILMV**VESSATSVEA**ALAQVR<br>RLKAAGAKRIAFDSGDVAHLKAAMEAGADILM**VAERHGLD**VALAQIRELKAAGAKEIAFASFDPDHL<br>LRAREEGADILVV**AGATDP**ARALATVRYLRAW |
| **KempTIM4h** | 233 | 24.9 | HHHHHHENLYFQSRIAYA**ALASDL**DSLVEALKLGADILLV**ILMADAAAVRAYAEMLHL**N**GDPRSVAE**<br>**LEALLRPTDLD**DALAAVRELKALGAKEIAFLSHDVDHLIRAMEAGADILMV**LESSATSVEA**ALAQVR<br>RLKAAGAKRISFDSGDVAHLKAAMEAGADILA**VAERHGLD**VALAQIRELKAAGAKEIAFASFDPDHL<br>LRAREEGADILVV**AGATDP**ARALATVRYLRAW |
| **KempTIM4-gsg** [c] | 236 | 25.1 | HHHHHHENLYFQSGSGRIAYM**ALASDL**DSLVEALKLGADILMV**ALMADAAAVRAMAEGLHA**N**GDPRS**<br>**VAELEALLRPTDLD**DALAAVRELKALGAKEIAFMSHDVDHLIRAMEAGADILMV**LESSATSVEA**ALA<br>QVRRLKAAGAKRISFGSGDVAHLKAAMEAGADILD**VLERHGLD**VALAQIRELKAAGAKEIAFASLDP<br>DHLLRAREEGADILVV**FGATDP**ARALATVRYLRAW |

[a] Molecular weight
[b] Catalytic and lid residues are underlined and bolded, respectively.
[c] This enzyme contains an additional GSG linker between the His-tag and first TIM barrel residue.

**Supplementary Table 2.** Purification yields

| Enzyme | Yield (mg $L^{-1}$) [a] |
|---|---|
| **NT6-CP1** | – |
| **NT6-CP2** | 70 ± 10 |
| **7MCD** | 31 ± 3 |
| **Variant 1** | – |
| **Variant 2** | – |
| **Variant 3** | – |
| **Variant 4** | 3 ± 1 |
| **Variant 5** | – |
| **Variant 6** | 12 ± 6 |
| **Variant 7** | 8 |
| **Variant 8** | 9 |
| **Variant 9** | 16 ± 7 |
| **KempTIM1** | 17 ± 9 |
| **KempTIM2** | – |
| **KempTIM3** | 59 |
| **KempTIM4** | 30 ± 10 |
| **KempTIM5** | 7 |
| **KempTIM4a** | 7 ± 5 |
| **KempTIM4b** | 30 ± 20 |
| **KempTIM4c** | 40 ± 10 |
| **KempTIM4d** | 30 ± 20 |
| **KempTIM4e** | 30 ± 20 |
| **KempTIM4f** | 40 ± 20 |
| **KempTIM4g** | 28 ± 1 |
| **KempTIM4h** | 30 ± 20 |

[a] Data represent the average measurements from 1 to 17 independent protein batches, with error bars indicating the standard deviation. NT6-CP1, Variants 1, 2, 3 and 5, and KempTIM2 could not be expressed solubly.

**Supplementary Table 3.** Theoretical and experimentally determined molecular weight (MW) using SEC-MALS

| Protein | Theoretical MW (kDa) | Molecular mass determined experimentally (kDa) |
|---|---|---|
| **NT6-CP2** | 22.1 | 21.2 ± 0.02 |
| **KempTIM1** | 25.2 | 24.7 ± 0.02 |
| **KempTIM1 D24A** | 25.2 | 24.6 ± 0.03 |
| **KempTIM1 N131A** | 25.2 | 24.6 ± 0.02 |
| **KempTIM4** | 25.0 | 23.5 ± 0.02 |
| **KempTIM4 D153A** | 25.0 | 23.5 ± 0.02 |
| **KempTIM4 N46A** | 25.0 | 23.5 ± 0.02 |
| **KempTIM4b** | 24.8 | 24.4 ± 0.02 |

**Supplementary Table 4.** Kinetic parameters of KempTIMs

| Enzyme | Base | H-bond donor | pH | Substrate | $k_{cat}$ ($s^{-1}$) | $K_M$ (mM) | $k_{cat}/K_M$ ($M^{-1}$ $s^{-1}$) |
|---|---|---|---|---|---|---|---|
| *CANVAS* | | | | | | | |
| KempTIM1 | D24 | N131 | 7 | 5NBZ | 4.2 ± 0.2 | 3.1 ± 0.2 | 1,400 ± 100 |
| | D24 | N131 | 10 | 5NBZ | 13.7 ± 0.5 | 0.66 ± 0.06 | 21,000 ± 2,000 |
| | D24 | N131 | 7 | 6NBZ | 5.6 ± 0.9 | 2.2 ± 0.5 | 2,500 ± 700 |
| | D24 | N131 | 10 | 6NBZ | 27 ± 3 | 0.6 ± 0.1 | 45,000 ± 9,000 |
| | D24A | N131 | 7 | 5NBZ | N.D. | N.D. | N.D. |
| | D24 | N131A | 7 | 5NBZ | 0.65 ± 0.02 | 2.6 ± 0.1 | 250 ± 10 |
| KempTIM3 | D153 | N46 | 7 | 5NBZ | 0.0176 ± 0.0004 | 2.11 ± 0.08 | 8.3 ± 0.4 |
| KempTIM4 | D153 | N46 | 7 | 5NBZ | N.D. | N.D. | 7.01 ± 0.08 |
| | D153 | N46 | 10 | 5NBZ | N.D. | N.D. | 18.6 ± 0.4 |
| | D153A | N46 | 7 | 5NBZ | N.D. | N.D. | N.D. |
| | D153 | N46A | 7 | 5NBZ | N.D. | N.D. | 6.5 ± 0.1 |
| KempTIM5 | D153 | N46 | 7 | 5NBZ | N.D. | N.D. | 1.60 ± 0.02 |
| Variant 4 | D153 | N46 | 7 | 5NBZ | 0.0066 ± 0.0006 | 0.6 ± 0.1 | 11 ± 3 |
| *Ensemble-based design* | | | | | | | |
| KempTIM4a | D153 | Q42 | 7 | 5NBZ | 0.55 ± 0.01 | 1.18 ± 0.06 | 470 ± 20 |
| KempTIM4b | D153 | Q42 | 7 | 5NBZ | 1.21 ± 0.04 | 0.39 ± 0.04 | 3,100 ± 300 |
| | D153 | Q42 | 10 | 5NBZ | 26.3 ± 0.9 | 0.83 ± 0.06 | 32,000 ± 3,000 |
| | D153 | Q42 | 7 | 6NBZ | 0.139 ± 0.008 | 0.22 ± 0.04 | 600 ± 100 |
| | D153 | Q42 | 10 | 6NBZ | 1.02 ± 0.02 | 0.27 ± 0.02 | 3,800 ± 300 |
| | D153A | Q42 | 7 | 5NBZ | N.D. | N.D. | N.D. |
| | D153 | Q42A | 7 | 5NBZ | 0.46 ± 0.03 | 0.7 ± 0.1 | 640 ± 90 |
| KempTIM4c | D5 | Q43 | 7 | 5NBZ | 0.18 ± 0.01 | 2.6 ± 0.2 | 69 ± 6 |
| KempTIM4d | D134 | N46 | 7 | 5NBZ | 0.61 ± 0.01 | 0.89 ± 0.04 | 690 ± 30 |
| KempTIM4e | D134 | N46 | 7 | 5NBZ | 0.15 ± 0.01 | 2.8 ± 0.2 | 54 ± 5 |
| KempTIM4f | D134 | N46 | 7 | 5NBZ | N.D. | N.D. | 19 ± 2 |
| KempTIM4g | D134 | N46 | 7 | 5NBZ | 0.42 ± 0.06 | 3.1 ± 0.7 | 140 ± 30 |
| KempTIM4h | D134 | N46 | 7 | 5NBZ | N.D. | N.D. | 9.7 ± 0.4 |

5NBZ and 6NBZ are 5-nitrobenzisoxazole and 6-nitrobenzisoxazole, respectively. $k_{cat}$ and $K_M$ were calculated by fitting the data to the Michaelis-Menten model $v_0 = k_{cat}[E_0][S]/(K_M+[S])$. Errors of nonlinear regression fitting, which represent the absolute measure of the typical distance that each data point falls from the regression line, are provided. N.D. indicates that individual parameters $K_M$ and $k_{cat}$ could not be determined accurately because saturation was not possible at the maximum substrate concentration tested (2 mM or 1 mM for 5NBZ or 6NBZ, respectively), which is the substrate's solubility limit. In those cases, catalytic efficiencies ($k_{cat}/K_M$) were calculated from the slope of the linear portion ($[S] \ll K_M$) of the Michaelis-Menten model ($v = (k_{cat}/K_M)[E_0][S]$). n = 1 independent protein batch for KempTIM3 and KempTIM5. n = 2 independent protein batches for KempTIM1 and all ensemble-based designs. n = 4 independent protein batches for KempTIM4 and variant 4. At least two technical replicates were performed for each independent batch. Kinetic assays were carried out in 50 mM sodium phosphate containing 100 mM NaCl and 10% MeOH (pH 7) or 50 mM CHES containing 100 mM NaCl and 10% MeOH (pH 10).

**Supplementary Table 5.** Kinetic parameters of most active KempTIMs at various pH

| Enzyme | pH | $k_{cat}$ ($s^{-1}$) | $K_M$ (mM) | $k_{cat}/K_M$ ($M^{-1} s^{-1}$) |
|---|---|---|---|---|
| **KempTIM1** | 7.0 | 4.2 ± 0.2 | 3.1 ± 0.2 | 1400 ± 100 |
| | 7.5 | N.D. | N.D. | 700 ± 50 |
| | 8.0 | N.D. | N.D. | 1700 ± 100 |
| | 8.5 | 4.4 ± 0.3 | 1.3 ± 0.2 | 3400 ± 500 |
| | 9.0 | 4.0 ± 0.1 | 0.37 ± 0.03 | 11,000 ± 1000 |
| | 9.5 | 12.3 ± 0.3 | 0.79 ± 0.04 | 15,600 ± 900 |
| | 10.0 | 13.7 ± 0.5 | 0.66 ± 0.06 | 21,000 ± 2000 |
| **KempTIM4b** | 7.0 | 1.21 ± 0.04 | 0.39 ± 0.04 | 3100 ± 300 |
| | 7.5 | 0.84 ± 0.03 | 0.42 ± 0.04 | 2000 ± 200 |
| | 8.0 | 2.14 ± 0.08 | 0.43 ± 0.05 | 5000 ± 600 |
| | 8.5 | 4.0 ± 0.1 | 0.43 ± 0.03 | 9300 ± 700 |
| | 9.0 | 13.9 ± 0.7 | 1.1 ± 0.1 | 13,000 ± 2000 |
| | 9.5 | 22 ± 1 | 1.1 ± 0.1 | 20,000 ± 3000 |
| | 10.0 | 26.3 ± 0.9 | 0.84 ± 0.06 | 32,000 ± 3000 |

$k_{cat}$ and $K_M$ using 5-nitrobenzisoxazole as substrate were calculated by fitting the data to the Michaelis-Menten model $v_0 = k_{cat}[E_0][S]/(K_M+[S])$. Errors of nonlinear regression fitting, which represent the absolute measure of the typical distance that each data point falls from the regression line, are provided. N.D. indicates that individual parameters $K_M$ and $k_{cat}$ could not be determined accurately because saturation was not possible at the maximum substrate concentration tested (2 mM), which is the substrate's solubility limit. In those cases, catalytic efficiencies ($k_{cat}/K_M$) were calculated from the slope of the linear portion ($[S] \ll K_M$) of the Michaelis-Menten model ($v = (k_{cat}/K_M)[E_0][S]$). n = 2 independent protein batches. At least two technical replicates were performed for each independent batch. Kinetic assays were carried out in 50 mM sodium phosphate containing 100 mM NaCl and 10% MeOH (pH 7.0), 50 mM Tris containing 100 mM NaCl and 10% MeOH (pH 7.5–8.5), or 50 mM CHES with 100 mM NaCl and 10% MeOH (pH 9.0–10.0).

**Supplementary Table 6.** Crystallographic data collection and refinement statistics

| | KempTIM1 + 6NBT | KempTIM1 apo | KempTIM4 apo |
|---|---|---|---|
| **PDB ID** | **9TZD** | **9U01** | **9QKX** |
| **Data collection** [a] | | | |
| Beamline | BESSY BL14.1 | BESSY BL14.1 | BESSY BL14.1 |
| Wavelength [Å] | 0.9184 | 0.9184 | 0.9184 |
| Space group | $P\ 2_1\ 2_1\ 2_1$ | $P\ 2_1\ 2_1\ 2_1$ | $P\ 3_1\ 2\ 1$ |
| Unit cell [Å, °] | a = 41.22 b = 67.27 c = 70.92<br>α = β = γ = 90 | a = 41.17 b = 66.69 c = 71.14<br>α = β = γ = 90 | a = b = 59.8 c = 120.69<br>α = β = 90 γ = 120 |
| Resolution range [Å] | 33.63-1.2<br>(1.23-1.2) | 33.35-1.25<br>(1.28-1.25) | 47.59-2.30<br>(2.53-2.30) |
| Unique reflections | 61955 (4030) | 54927 (3588) | 11469 (2772) |
| Multiplicity | 13.0 (12.4) | 12.6 (13.2) | 19.5 (20.4) |
| Completeness [%] | 99.22 (98.01) | 99.98 (99.94) | 98.65 (98.30) |
| *R-meas* [%] | 0.07119 (2.28) | 0.07025 (2.128) | 0.4812 (2.946) |
| $\langle I/\sigma I \rangle$ | 16.97 (1.09) | 14.09 (1.03) | 8.24 (1.58) |
| $CC_{1/2}$ | 1 (0.468) | 1 (0.573) | 0.995 (0.404) |
| $CC^*$ | 1 (0.798) | 1 (0.854) | 0.999 (0.759) |
| Wilson *B*-factor [$Å^2$] | 15.18 | 17.30 | 33.87 |
| **Refinement** | | | |
| $R_{work}/R_{free}$ [%] | 15.55 / 17.73 | 15.70 / 17.74 | 25.47 / 29.96 |
| No. of atoms (non-H) | | | |
| macromolecules | 1752 | 1754 | 1640 |
| ligands | 36 | 16 | 42 |
| solvent | 191 | 185 | 52 |
| RMSD from ideal geometry | | | |
| bonds [Å] | 0.007 | 0.007 | 0.001 |
| angles [°] | 1.03 | 0.91 | 0.310 |
| Ramachandran statistics | | | |
| favored [%] | 100.00 | 99.08 | 98.63 |
| outliers [%] | 0.00 | 0.00 | 0.00 |
| Clashscore | 5.24 | 0.83 | 4.96 |
| Average *B* [$Å^2$] | 23.02 | 27.50 | 42.19 |
| macromolecules | 21.72 | 26.15 | 41.79 |
| ligands | 30.25 | 38.16 | 59.28 |
| solvent | 33.56 | 39.30 | 41.22 |

[a] Statistics for the highest-resolution shell are shown in parentheses.

**Supplementary Table 7.** Geometric constraints used to define catalytic contacts during computational design

| Contact | Residue | Type | Atom 1 [a] | Atom 2 [a] | Atom 3 [a] | Atom 4 [a] | Min [b] | Max [b] |
|---|---|---|---|---|---|---|---|---|
| **Base** | **Asp** | Distance | OD1 or OD2 | **H3** | | | 1.0 | 1.6 |
| | | Angle | CG | OD1 or OD2 | **H3** | | 109 | 131 |
| | | Angle | OD1 or OD2 | **H3** | **C3** | | 159 | 180 |
| | | Torsion | CB | CG | OD1 or OD2 | **H3** | –21, 159 | 21, 201 |
| **H-bond donor** | **Asn** | Distance | 1HD2 or 2HD2 | **O1** | | | 1.2 | 2.3 |
| | | Angle | ND2 | 1HD2 or 2HD2 | **O1** | | 145 | 157 |
| | | Angle | 1HD2 or 2HD2 | **O1** | **N2** | | 120 | 140 |
| | | Torsion | 1HD2 or 2HD2 | **O1** | **N2** | **C3** | –20, 160 | 20, 200 |
| | **Gln** | Distance | 1HE2 or 2HE2 | **O1** | | | 1.2 | 2.3 |
| | | Angle | NE2 | 1HE2 or 2HE2 | **O1** | | 145 | 157 |
| | | Angle | 1HE2 or 2HE2 | **O1** | **N2** | | 120 | 140 |
| | | Torsion | 1HE2 or 2HE2 | **O1** | **N2** | **C3** | –20, 160 | 20, 200 |

[a] Atoms in bold are from the transition state. All other atoms are from the catalytic residues.
[b] Distance measurements given in Å, all others in degrees.

**Supplementary Table 8.** Amino-acid positions for theozyme placement

| Input structure | Residues for catalytic base placement [a] | Residues mutated to alanine [a] |
|---|---|---|
| **7MCD** | I4, I31, I50, I77, I96, I123, I142, I169 | N6, I31, N52, I77, N98, I123, N144, I169 |
| **NT6-CP1** | R5, I24, R51, I70, R97, I116, R143, I162 | A3, R5, I24, D26, A49, R51, I70, D72, A95, R97, I116, D118, A141, R143, I162, D164 |
| **NT6-CP2** | R5, I24, R51, I70, R97, I116, R143, I162 | A3, R5, I24, D26, A49, R51, I70, D72, A95, R97, I116, D118, A141, R143, I162, D164 |

[a] Residue numbering based on input structure

**Supplementary Table 9.** Geometric definitions for generation of transition-state poses off the side chain of catalytic base and H-bond donor

| Catalytic residue | Type | Atom 1 [a] | Atom 2 [a] | Atom 3 [a] | Atom 4 [a] | Values [b] |
|---|---|---|---|---|---|---|
| **Asp** | Distance | OD1 or OD2 | **H3** | | | 1.0, 1.2, 1.5 |
| | Angle | CG | OD1 or OD2 | **H3** | | 112, 117, 122 |
| | Angle | OD1 or OD2 | **H3** | **C3** | | 159, 164, 169, 174, 179 |
| | Torsion | CB | CG | OD1 or OD2 | **H3** | 0, 5, 10, 170, 175, 180 |
| | Torsion | CG | OD1 or OD2 | **H3** | **C3** | 170, 175, 180, 185, 190 |
| | Torsion | OD1 or OD2 | **H3** | **C3** | **N2** | 0, 5, 170, 175, 180 |
| **Asn** | Distance | 1HD2 or 2HD2 | **O1** | | | 1.2, 1.5, 1.8, 2.1, 2.3 |
| | Angle | ND2 | 1HD2 or 2HD2 | **O1** | | 145, 151, 157 |
| | Angle | 1HD2 or 1HD2 | **O1** | **N2** | | 120, 135, 140 |
| | Torsion | CG | ND2 | 1HD1 or 2HD2 | **O1** | 120, 135, 140 |
| | Torsion | ND2 | 1HD2 or 2HD2 | **O1** | **N2** | –121, –116, –111 |
| | Torsion | 1HD2 or 2HD2 | **O1** | **N2** | **C3** | 160, 180, 200 |
| **Gln** | Distance | 1HE2 or 2HE2 | **O1** | | | 1.2, 1.5, 1.8, 2.1, 2.3 |
| | Angle | NE2 | 1HE2 or 2HE2 | **O1** | | 145, 151, 157 |
| | Angle | 1HE2 or 2HE2 | **O1** | **N2** | | 120, 135, 140 |
| | Torsion | CD | NE2 | 1HE2 or 2HE2 | **O1** | 120, 135, 140 |
| | Torsion | NE2 | 1HE2 or 2HE2 | **O1** | **N2** | –121, –116, –111 |
| | Torsion | 1HE2 or 2HE2 | **O1** | **N2** | **C3** | 160, 180, 200 |

[a] Atoms in bold are from the transition state. All other atoms are from the catalytic residue.
[b] Distance measurements given in Å, all others are in degrees.

**Supplementary Table 10.** Amino-acid positions optimized during active-site repacking of KempTIM1

| Designed positions | Allowed amino acids [a] |
|---|---|
| **3** | MET, ILE, LEU, PHE, **ALA**, VAL |
| **5** | **ALA**, ILE, LEU, VAL, PHE |
| **49** | MET, ILE, **LEU**, PHE, ALA, VAL |
| **51** | **ALA** |
| **70** | **ALA** |
| **95** | MET, ILE, LEU, PHE, **ALA**, VAL |
| **97** | **GLY** |
| **116** | **GLY** |
| **118** | **ALA** |
| **127** | VAL, ILE, **LEU**, TYR, PHE, TRP, ALA |
| **173** | **MET**, ILE, LEU, PHE, ALA, VAL |
| **175** | **ALA**, ILE, LEU, VAL, PHE |
| **194** | ALA, ILE, **LEU**, VAL, PHE |
| **196** | **ALA**, ILE, LEU, VAL, PHE |

[a] Residues in bold are mutations found at those positions in the KempTIM1 sequence.

**Supplementary Table 11.** Design filtering

| Enzyme | Base dihedral (°) | H-bond dihedral (°) | Energy (kcal mol$^{-1}$) [a] | SASA (Å$^2$) [b] | # residues not preorganized [c] | Tunnel present [d] |
|---|---|---|---|---|---|---|
| **Variant 1** | –42.5 | 25.7 | –39.4 | 174 | 5 | Yes |
| **Variant 2** | –7.3 | 25.7 | –36.3 | 188 | 6 | Yes |
| **Variant 3** | 27.6 | –156.2 | –10.4 | 107 | 6 | Yes |
| **Variant 4** | 27.6 | –156.2 | –2.3 | 107 | 5 | Yes |
| **Variant 5** | 27.6 | –156.2 | –10.4 | 107 | 6 | Yes |
| **Variant 6** | –1 | 28.8 | –115.54 | 87 | 2 | Yes |
| **Variant 7** | –1 | 28.8 | –111.84 | 87 | 2 | Yes |
| **Variant 8** | –0.7 | 44.2 | –108.87 | 100 | 3 | Yes |
| **Variant 9** | –13.3 | 29.6 | –107.41 | 91 | 2 | Yes |

[a] Energies are calculated by subtracting the energy of an all-Gly structure as described in the methods under *Active-site repacking*.
[b] Solvent-accessible surface area of the transition state in the design model.
[c] Preorganized residues are those predicted to adopt the same rotamer in the presence and absence of the transition state. Variant 1–2, Variant 3–5, and Variant 6–9 had 14, 17, and 18 residues that were optimized during active-site repacking, respectively.
[d] The presence of a tunnel was determined by Caver 3.0 using a minimum cutoff bottleneck radius of 0.9 Å.

**Supplementary Table 12.** Ensemble refinement of KempTIM4

| | MD sampling parameters | | | Before ensemble refinement | | After ensemble refinement | | |
|---|---|---|---|---|---|---|---|---|
| **Ensemble** | $p_{TLS}$ | $\tau_x$ | $w_{x\text{-}ray}$ | $R_{work}$ | $R_{free}$ | $R_{work}$ | $R_{free}$ | **No. structures** |
| **1** | 0.8 | 1.5 | 5 | 0.2547 | 0.2996 | 0.2343 | 0.2782 | 20 |
| **2** | 0.8 | 2.0 | 10 | 0.2547 | 0.2996 | 0.2165 | 0.2808 | 40 |

**Supplementary Table 13.** Amino-acid positions for theozyme placement during ensemble-based design

| Input structure | Residues for catalytic base placement | Residues for catalytic H-bond donor placement | Residues mutated to alanine |
|---|---|---|---|
| **KempTIM4 crystallographic ensemble** | M5, M26, M85, M104, G134, D153, A181, V200 | M5, M26, M39, G42, L43, A45, N46, M85, F202 | A3, M5, M26, A28, A38, M39, G42, L43, A45, N46, A83, M85, M104, L106, S108, S132, G134, D153, L155, E156, R157, A179, A181, L183, V200, F202 |

**Supplementary Table 14.** Triad model filtering during ensemble-based design

| Metric | Residue | Type | Atom 1[a] | Atom 2[a] | Atom 3[a] | Atom 4[a] | Min[b] | Max[b] |
|---|---|---|---|---|---|---|---|---|
| **Base** | **Asp** | Distance | OD1 or OD2 | **H3** | | | | 1.6 |
| | | Angle | CG | OD1 or OD2 | **H3** | | 92 | 132 |
| | | Angle | OD1 or OD2 | **H3** | **C3** | | 139 | 179 |
| | | Torsion | CB | CG | OD1 or OD2 | **H3** | –20, 160 | 20, 200 |
| | | Torsion | CG | OD1 or OD2 | **H3** | **C3** | –20, 160 | 20, 200 |
| | | Torsion | OD1 or OD2 | **H3** | **C3** | **N2** | –30, 150 | 30, 210 |
| **H-bond donor** | **Asn** | Distance | 1HD2 or 2HD2 | **O1** | | | | 2.3 |
| | | Angle | ND2 | 1HD2 or 2HD2 | **O1** | | 131 | 171 |
| | | Angle | 1HD2 or 2HD2 | **O1** | **N2** | | 110 | 150 |
| | | Torsion | CB | CG | 1HD2 or 2HD2 | **O1** | –20, 160 | 20, 200 |
| | | Torsion | CG | 1HD2 or 2HD2 | **O1** | **N2** | –20, 160 | 20, 200 |
| | | Torsion | 1HD2 or 2HD2 | **O1** | **N2** | **C3** | –20, 160 | 20, 200 |
| | **Gln** | Distance | 1HE2 or 2HE2 | **O1** | | | | 2.3 |
| | | Angle | NE2 | 1HE2 or 2HE2 | **O1** | | 131 | 171 |
| | | Angle | 1HE2 or 2HE2 | **O1** | **N2** | | 110 | 150 |
| | | Torsion | CG | CD | 1HE2 or 2HE2 | **O1** | –20, 160 | 20, 200 |
| | | Torsion | CD | 1HE2 or 2HE2 | **O1** | **N2** | –20, 160 | 20, 200 |
| | | Torsion | 1HE2 or 2HE2 | **O1** | **N2** | **C3** | –20, 160 | 20, 200 |
| **Bottleneck radius[c]** | | Radius | | | | | 1.0 | |

[a] Atoms in bold are from the transition state. All other atoms are from the catalytic residues.
[b] Distance and radius measurements given in Å, all others in degrees.
[c] Determined with Caver 3.0 using a minimum cutoff bottleneck radius of 0.9 Å, only for predictions containing a tunnel

**Supplementary Table 15.** Filtering using Boltz-2 models during ensemble-based design

| Metric | Type | Atom 1 [a] | Atom 2 [a] | Min [b] | Max [b] |
|---|---|---|---|---|---|
| **Base distance** | Distance | OD1 or OD2 | **N3** | – | 3.3 |
| **H-bond donor** | Distance | ND2 or NE2 | **N1** | – | 3.3 |
| **Triad vs Boltz-2 transition-state analogue RMSD** [c] | RMSD | – | – | – | 1.0 |
| **Bound vs unbound active-site RMSD** [d] | RMSD | – | – | – | 1.0 |
| **Bottleneck radius** [e] | Radius | – | – | 1.2 | – |
| **Tunnel length** [e] | Distance | – | – | – | 15 |

[a] Atoms in bold are from the transition-state analogue. All other atoms are from the catalytic residues.
[b] All measurements given in Å.
[c] RMSD between the atoms of the benzene rings of the transition-state analogue in the Boltz2 model and transition-state in the Triad model.
[d] RMSD between the sidechains of active site residues in the Boltz2 transition-state analogue-bound and -unbound models
[e] Determined with Caver 3.0 using a minimum cutoff bottleneck radius of 0.9 Å, only for predictions containing a tunnel

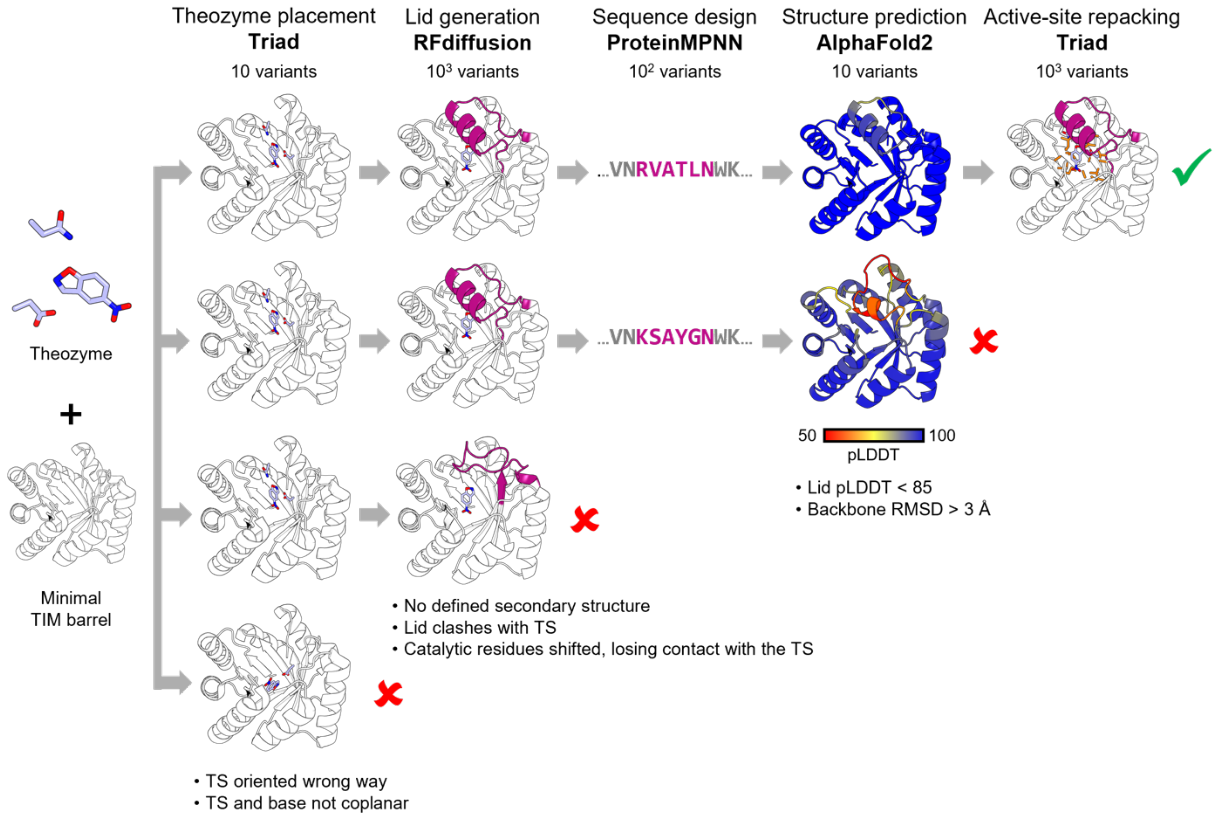


**Supplementary Figure 1. Detailed CANVAS workflow.** Step 1: A catalytic residue from the theozyme is placed onto a minimal TIM barrel scaffold, and the transition state (TS) is constructed from its side chain using Triad. The second catalytic residue is built from the TS, positioning its α-carbon in the empty space above the TIM barrel catalytic face. Additional catalytic residues can be added inside the barrel or the empty space using similar steps. Step 2: A lid composed of protein fragments of desired length and secondary structure is generated with RFdiffusion to anchor catalytic residues located above the TIM barrel face. Step 3: Lid sequences are designed with ProteinMPNN while maintaining catalytic residue identities. Step 4: Designed structures are predicted with AlphaFold2. Step 5: The amino acid sequence and rotameric configuration of the active site is optimized with Triad using the AlphaFold2 model as template to maximize TS packing and catalytic contact geometry. Resulting sequences are then filtered using key enzyme design criteria. At each step, structures failing filtering criteria are rejected. Approximate number of structures/sequences generated at each step are indicated as variants.

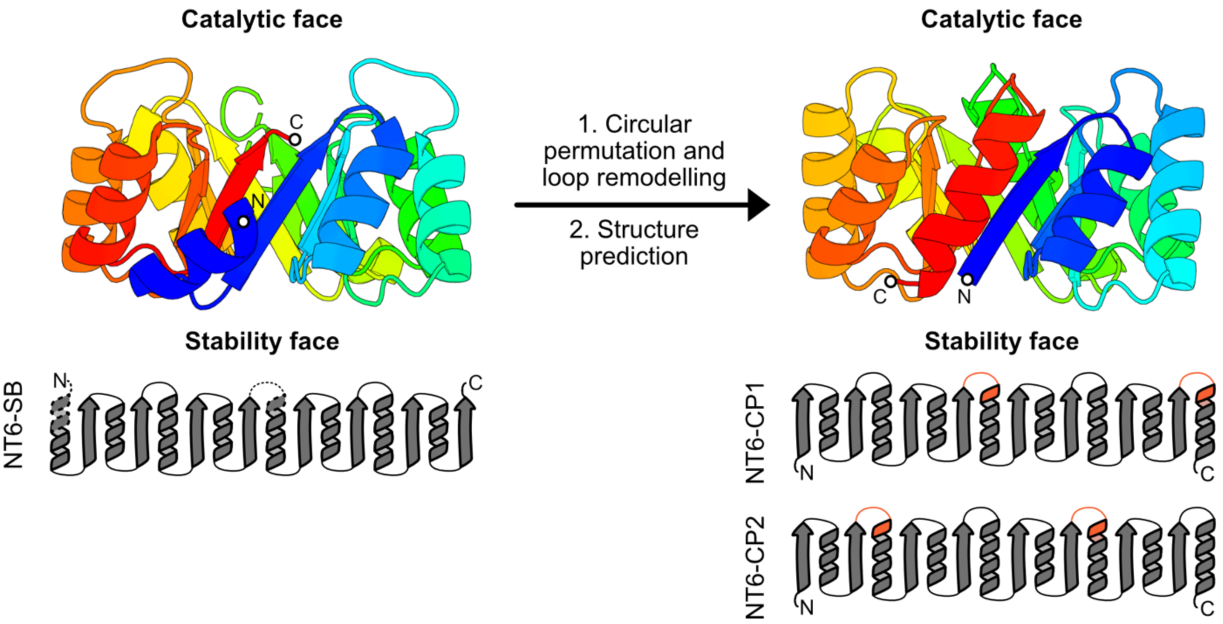


**Supplementary Figure 2. Circular permutation of *de novo* TIM barrel NT6-SB (PDB ID: 7OSV).** Circular permutation and loop remodeling were used to relocate the N- and C-termini of NT6-SB (left) from the catalytic face to the stability face. After structure prediction with AlphaFold2, variants NT6-CP1 and NT6-CP2 (right) were generated. Protein structures are colored from blue (N-terminus) to red (C-terminus). Topology diagrams show the secondary structural elements of TIM barrels. Dashed parts of NT6-SB indicate all non-resolved regions of the crystal structure. Orange regions in the circularly permuted TIM barrels represent the remodelled or inserted loops required for circular permutation.

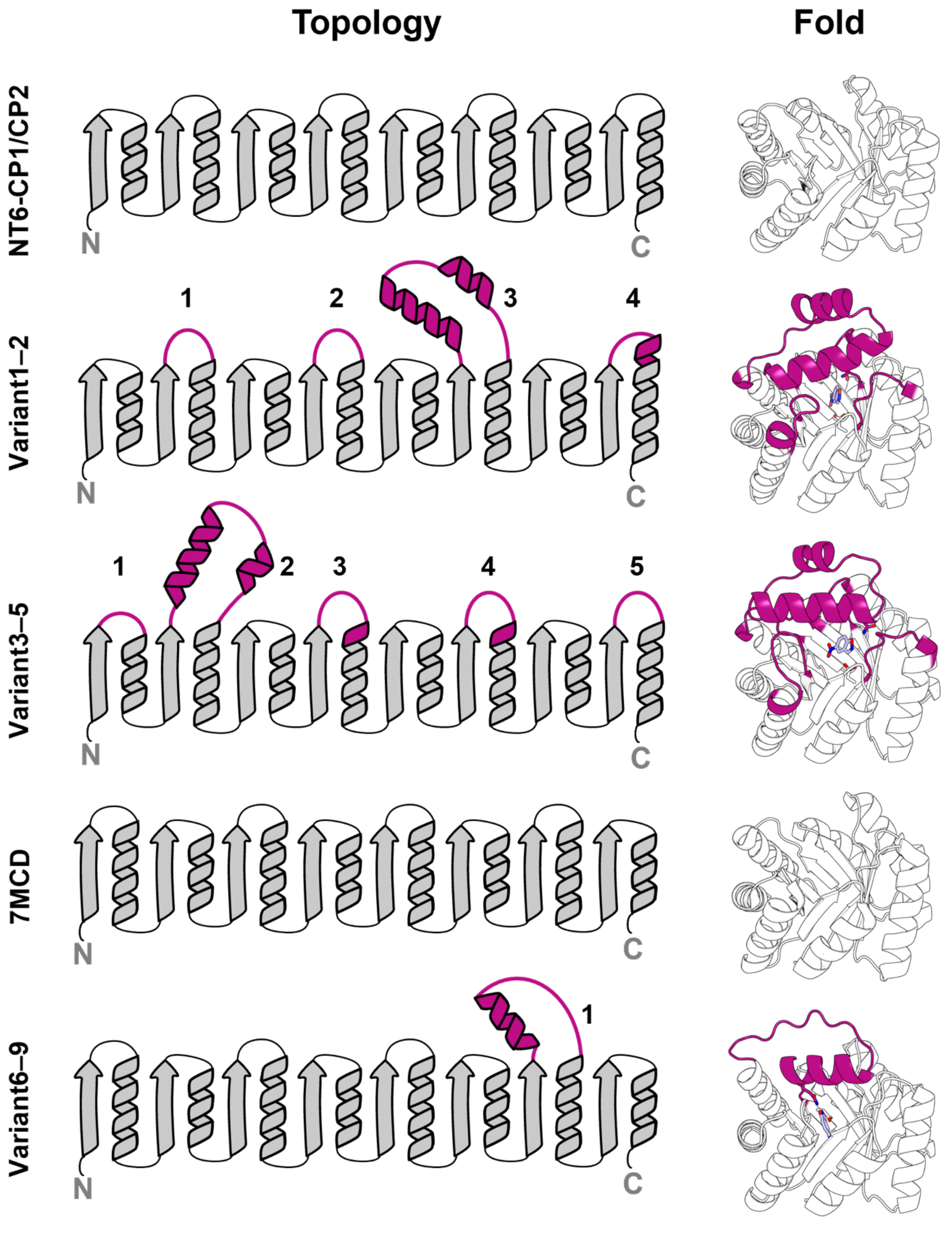


**Supplementary Figure 3. Structure of TIM barrels.** The topology and fold of various TIM barrels are shown. The designed lids (magenta) of Variant 1–2 and Variant 3–5 consist of four or five inserted fragments, including a long helix-turn-helix and three or four extended loops. The lids of Variant 6–9 contain a single inserted helix-loop fragment.

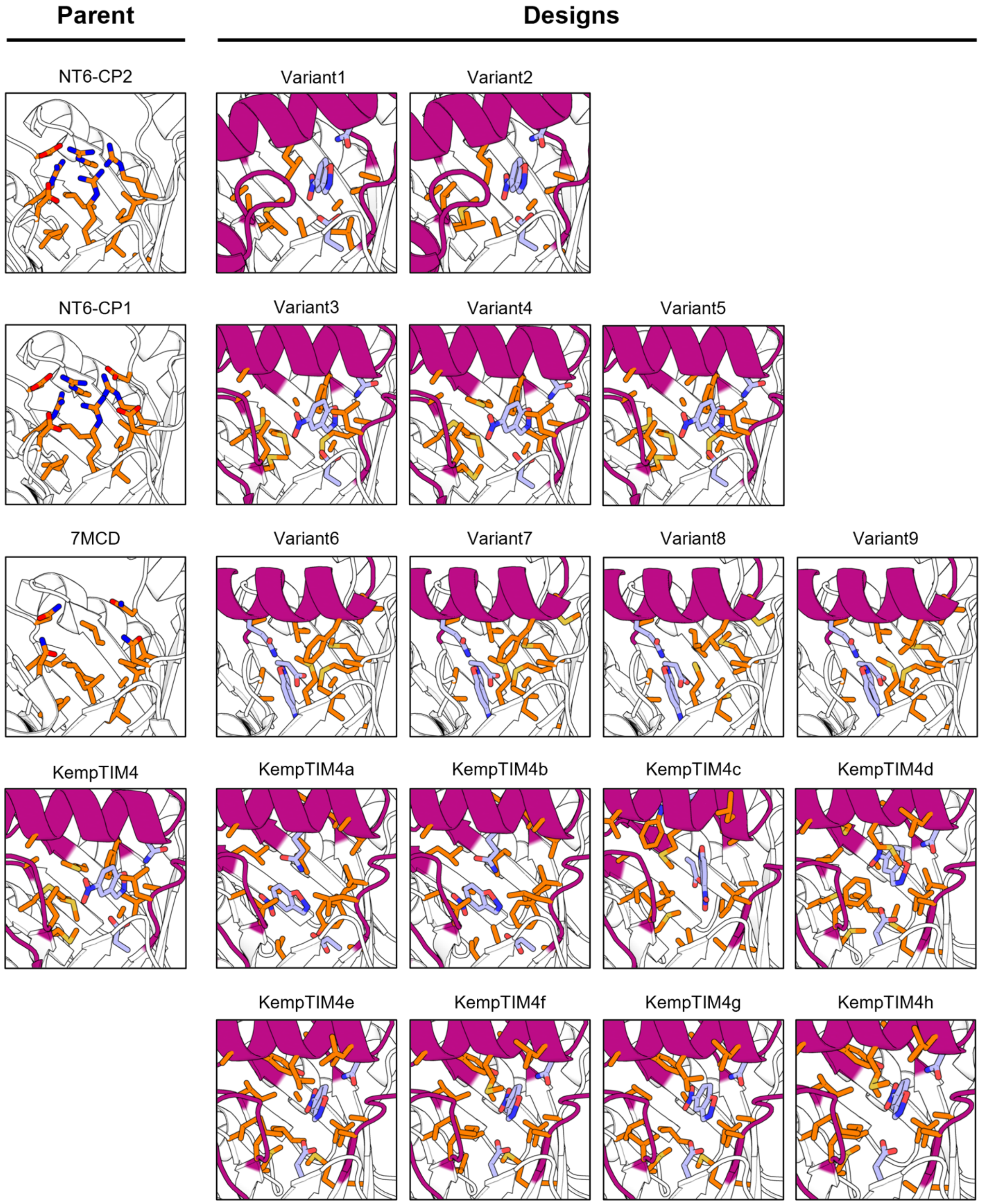


**Supplementary Figure 4. Active site configurations of designed enzymes.** Residues optimized by Triad during active-site repacking of variants, along with the corresponding residues on the parent minimal TIM barrel, are shown as orange sticks. Lids introduced by RFdiffusion are shown in magenta. The catalytic base, catalytic hydrogen-bond donor, and transition state are depicted as light blue sticks.

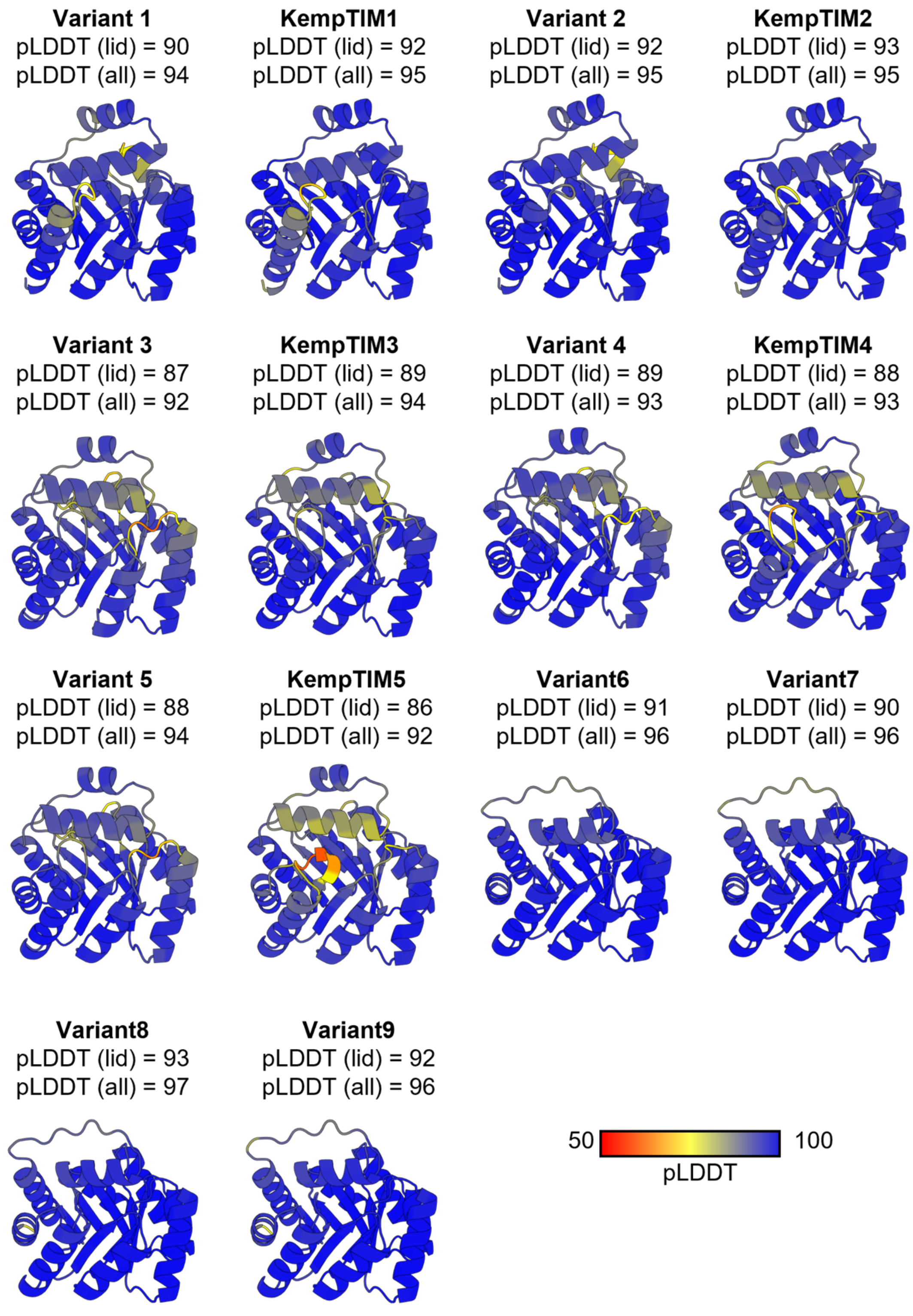


**Supplementary Figure 5. AlphaFold2 models.** Structures of variants are coloured by pLDDT, with the average pLDDT of lid residues and the entire structure listed above each variant.

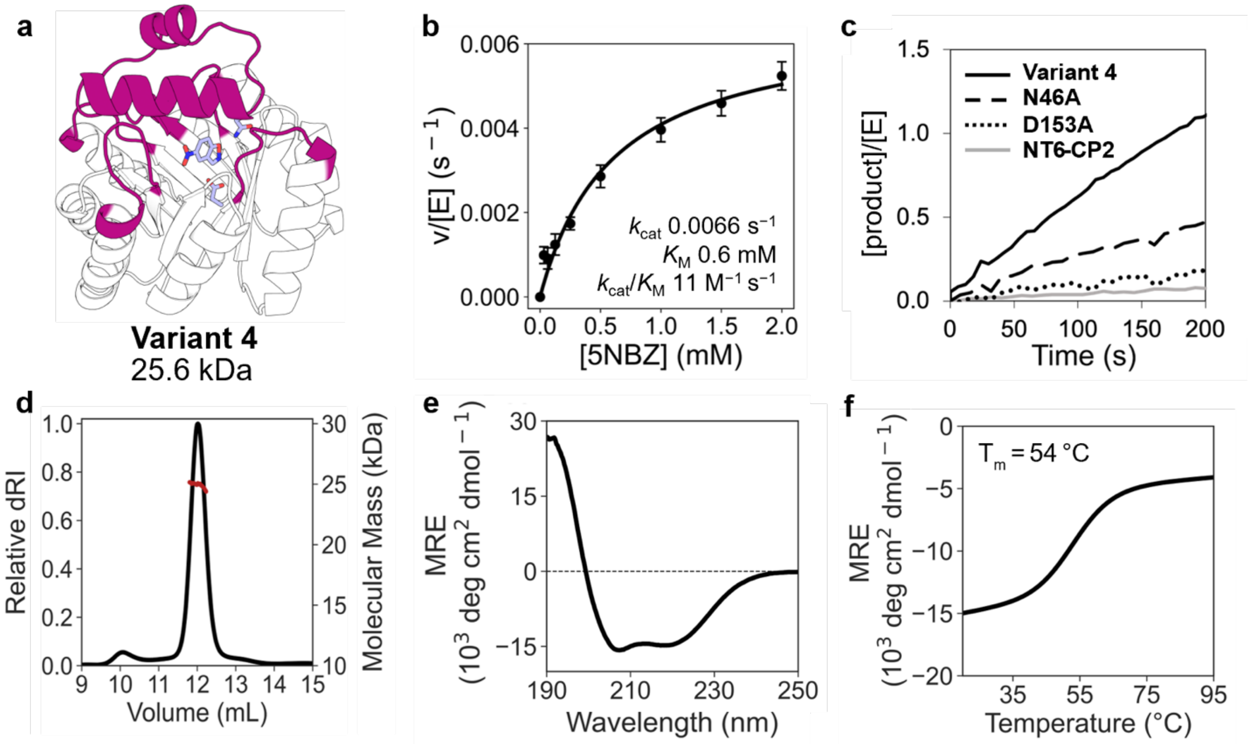


**Supplementary Figure 6. Structural and kinetic characterization of variant 4.** (a) Computational model of variant 4 with the designed lid shown in magenta and the theozyme as sticks. (b) Michaelis–Menten plot of normalized initial rates as a function of 5-nitrobenzisoxazole (5NBZ) concentrations. Data represent the average of 15 replicate measurements from five independent protein batches (mean ± SEM). (c) Reaction progress curves ([5NBZ] = 2 mM) demonstrate the contributions of designed catalytic residues D153 and N46 to catalysis. (d) SEC-MALS indicates that variant 4 is predominantly monomeric with a minor tendency to form dimers. dRI: differential refractive index. (e) CD spectrum reveals a mixed αβ signal characteristic of TIM barrels. MRE: mean residue ellipticity. (f) Melting curves demonstrate that variant 4 is substantially destabilized compared to NT6-CP2 ($T_m$ = 89 °C). $T_m$: melting temperature.

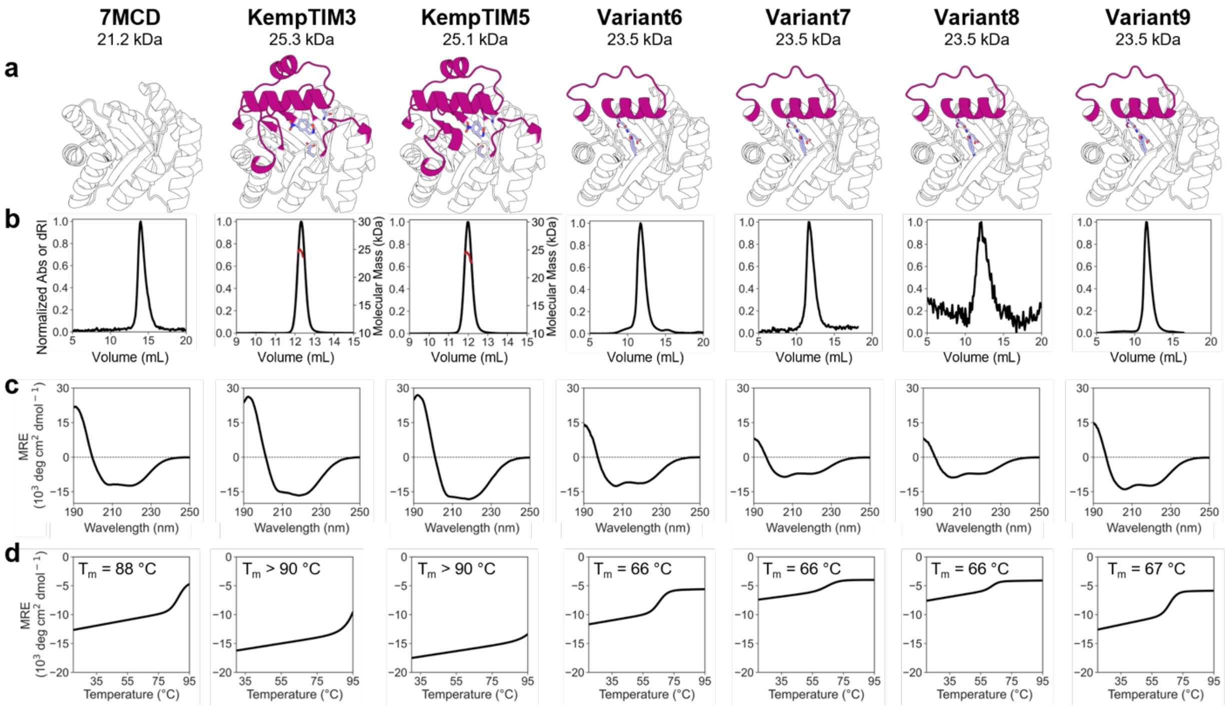


**Supplementary Figure 7. Structural characterization.** (a) Computational models of minimal TIM-barrel 7MCD and variants with the designed lid shown in magenta and the theozyme as sticks. (b) SEC-MALS and SEC indicate that the proteins are predominantly monomeric. Abs: absorbance. dRI: differential refractive index. (c) Circular dichroism spectra reveal a mixed αβ signal characteristic of TIM barrels. MRE: mean residue ellipticity. (d) Melting curves demonstrate that Variants 6–9 are substantially destabilized compared to their parent 7MCD. $T_m$: melting temperature.

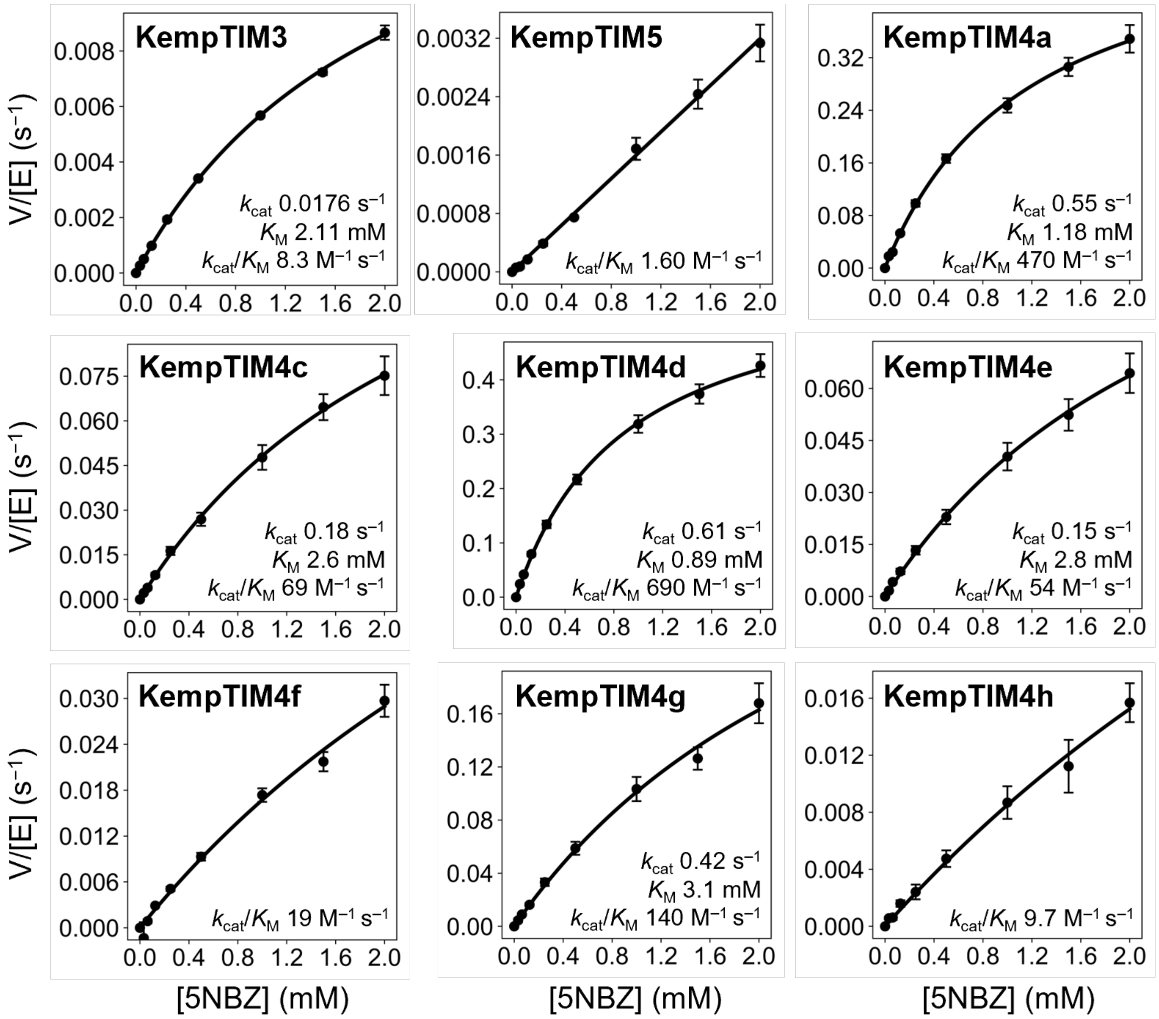


**Supplementary Figure 8. Kinetic characterization of KempTIMs.** Kinetic assays were carried out in 50 mM sodium phosphate containing 100 mM NaCl and 10% MeOH (pH 7), Michaelis-Menten plots showing normalized initial rates as a function of 5-nitrobenzisoxazole (5NBZ) concentration. Data represent the mean ± SEM of ≥ 3 replicate measurements from ≥ 1 independent protein batches.

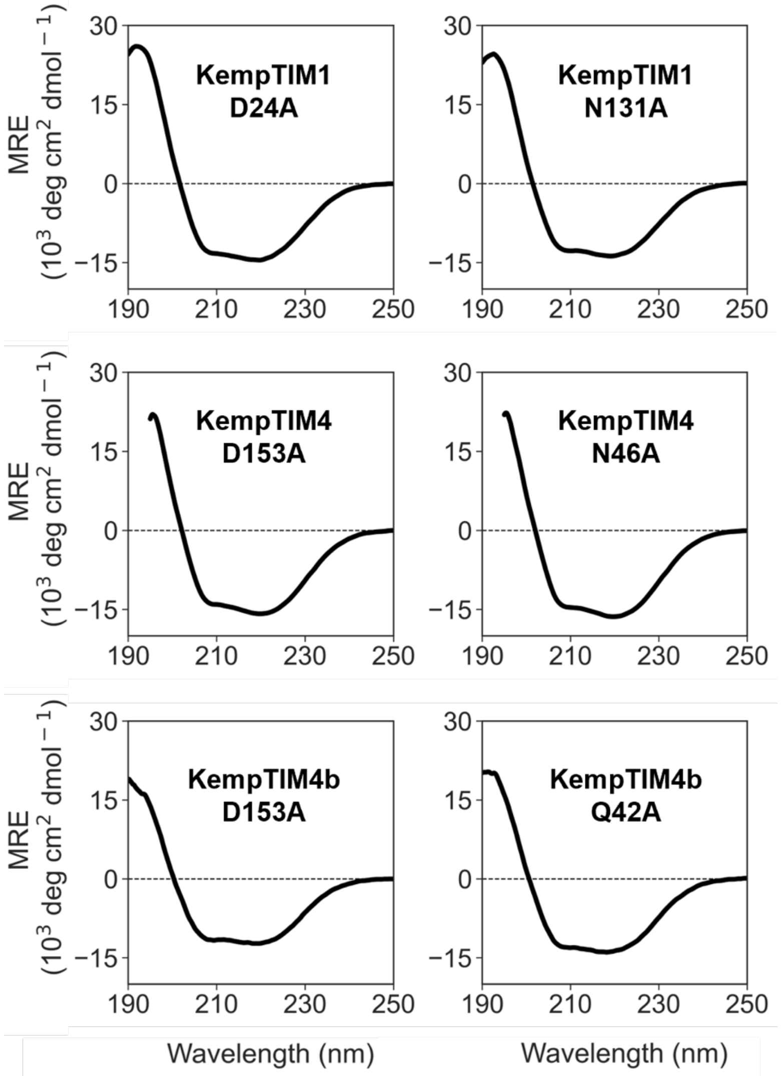


**Supplementary Figure 9. Structural characterization of catalytic knockout mutants.** Circular dichroism spectra demonstrate that mutation of catalytic residues does not change the structure. MRE: mean residue ellipticity.

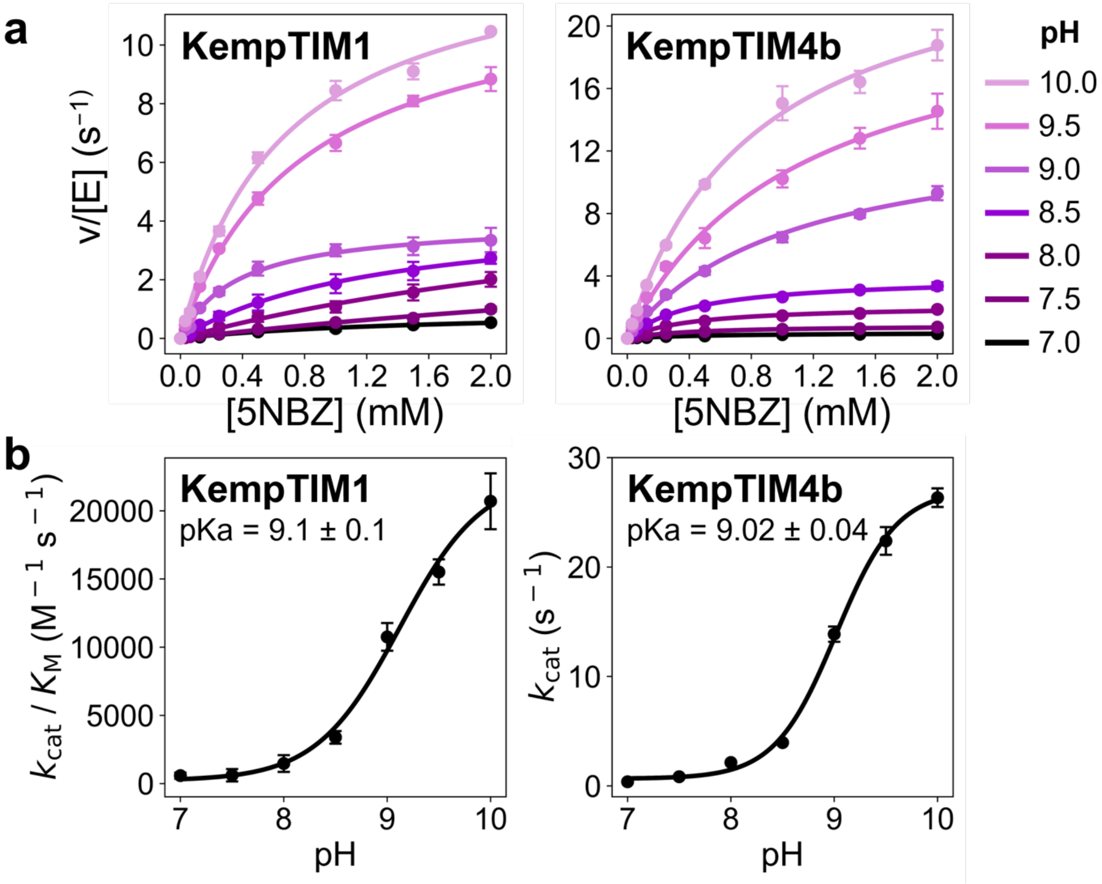


**Supplementary Figure 10. Activity of KempTIM1 and KempTIM4b increases under alkaline conditions.** Kinetic assays were performed in 50 mM Tris buffer containing 100 mM NaCl and 10% MeOH (pH 7.0–8.5) or in 50 mM CHES buffer containing 100 mM NaCl and 10% MeOH (pH 9.0–10.0). Data represent mean ± SEM from ≥ 4 replicate measurements across ≥ 2 independent protein batches. (a) Michaelis–Menten plots at varying pH showing normalized initial rates as a function of 5-nitrobenzisoxazole (5NBZ) concentration. (b) pH–rate profiles reveal a single inflection near pH 9, with maximal enzymatic activity observed at pH ≥ 10.

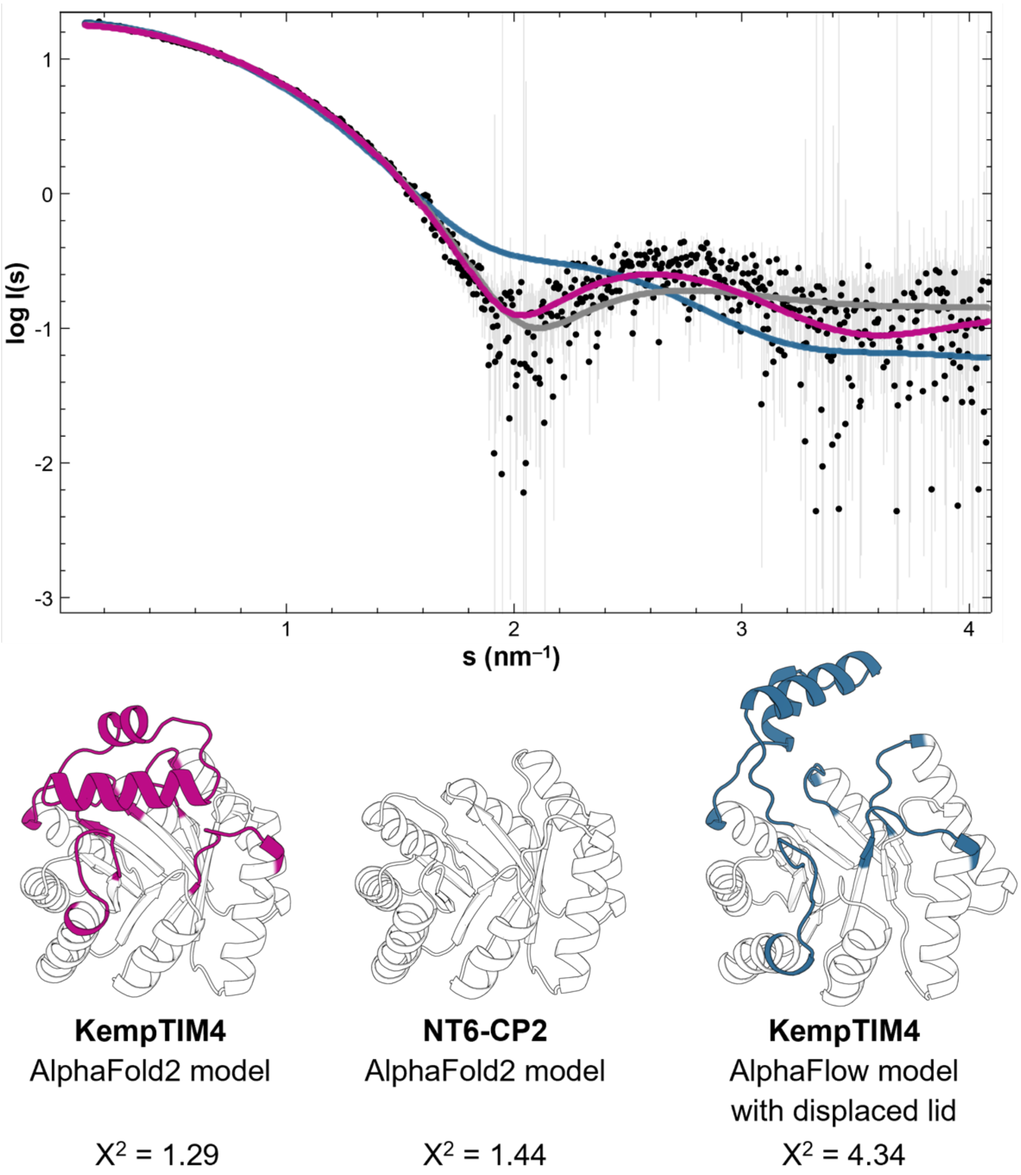


**Supplementary Figure 11. Comparison of KempTIM4 experimental SAXS data with scattering calculated from various similar size TIM barrels.** Experimental scattering data for KempTIM4 (black) align more closely with the calculated scattering curve from the KempTIM4 AlphaFold2 model (magenta) than with models of the minimal TIM barrel NT6-CP2 (grey) or a KempTIM4 model with improperly positioned lid generated using AlphaFlow (blue).

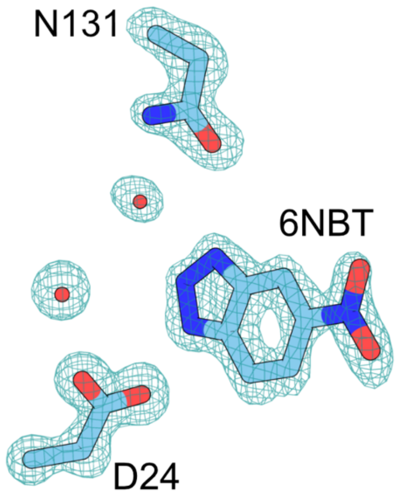


**Supplementary Figure 12. Polder map of 6NBT, catalytic residues D24 and N131 and interacting water molecules.** Polder map contoured at ±6σ. Calculated map shows a well-defined density for all displayed atoms.

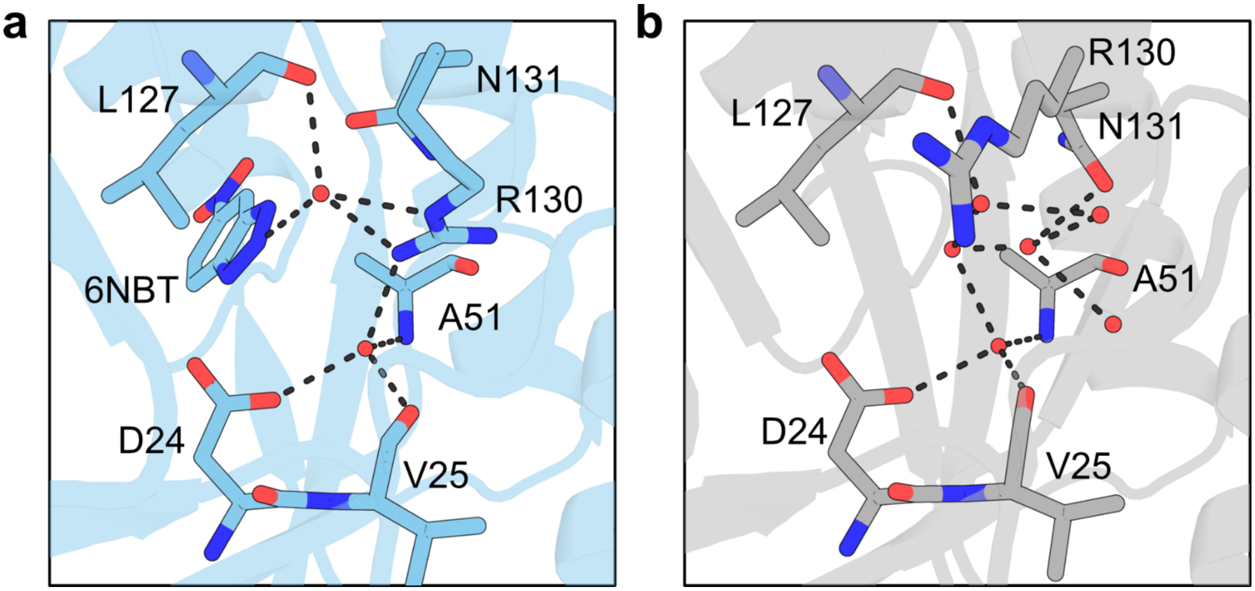


**Supplementary Figure 13. Water networks in the active site of KempTIM1.** (a) In the co-crystal structure, two ordered water molecules are observed: one hydrogen bonds to the catalytic base D24 and the other to the transition-state analogue 6NBT. Both waters also interact with R130, which adopts a conformation that positions its guanidinium group within the active-site pocket. (b) In the apo structure, six ordered water molecules are present owing to the absence of bound ligand and a rotameric change in R130 that displaces its side chain away from the active-site pocket. Several of these water molecules occupy positions similar to that of the R130 guanidinium group in the co-crystal structure.

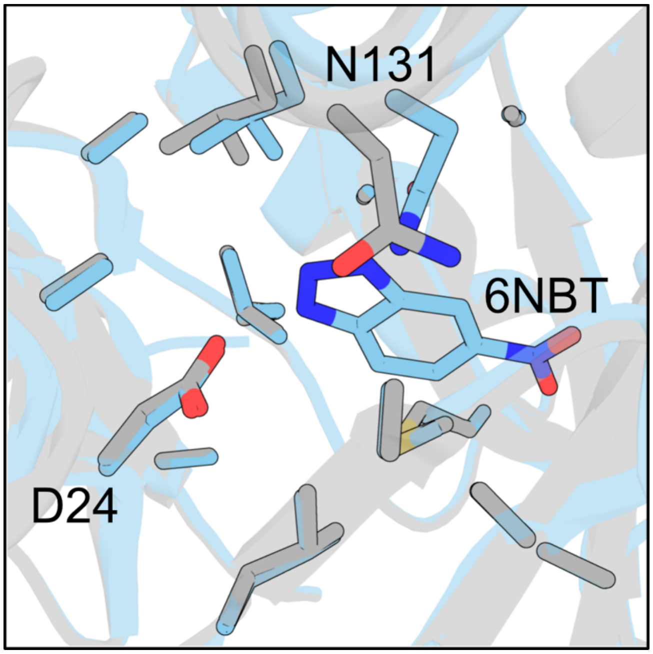


**Supplementary Figure 14. Transition-state analogue binding induces lid rearrangement in KempTIM1.** Comparison of the active sites in the co-crystal (blue) and apo (grey) structures shows that residues within the minimal TIM barrel adopt identical side-chain rotamers in both states, indicating high active-site preorganization. By contrast, the lid undergoes a subtle but concerted shift upon ligand binding, resulting in altered Cα positions for the catalytic H-bond donor N131 and a corresponding rotameric change. This lid displacement indicates localized structural rearrangements associated with transition-state analogue binding.

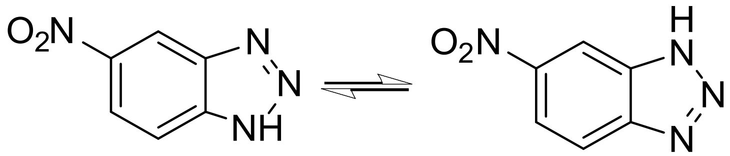


**Supplementary Figure 15. Tautomeric forms of 6-nitrobenzotriazole.** The 6-nitrobenzotriazole transition-state analogue tautomerizes in water, which allows it to bind to enzymes that react with either 5- or 6-nitrobenzisoxazole.

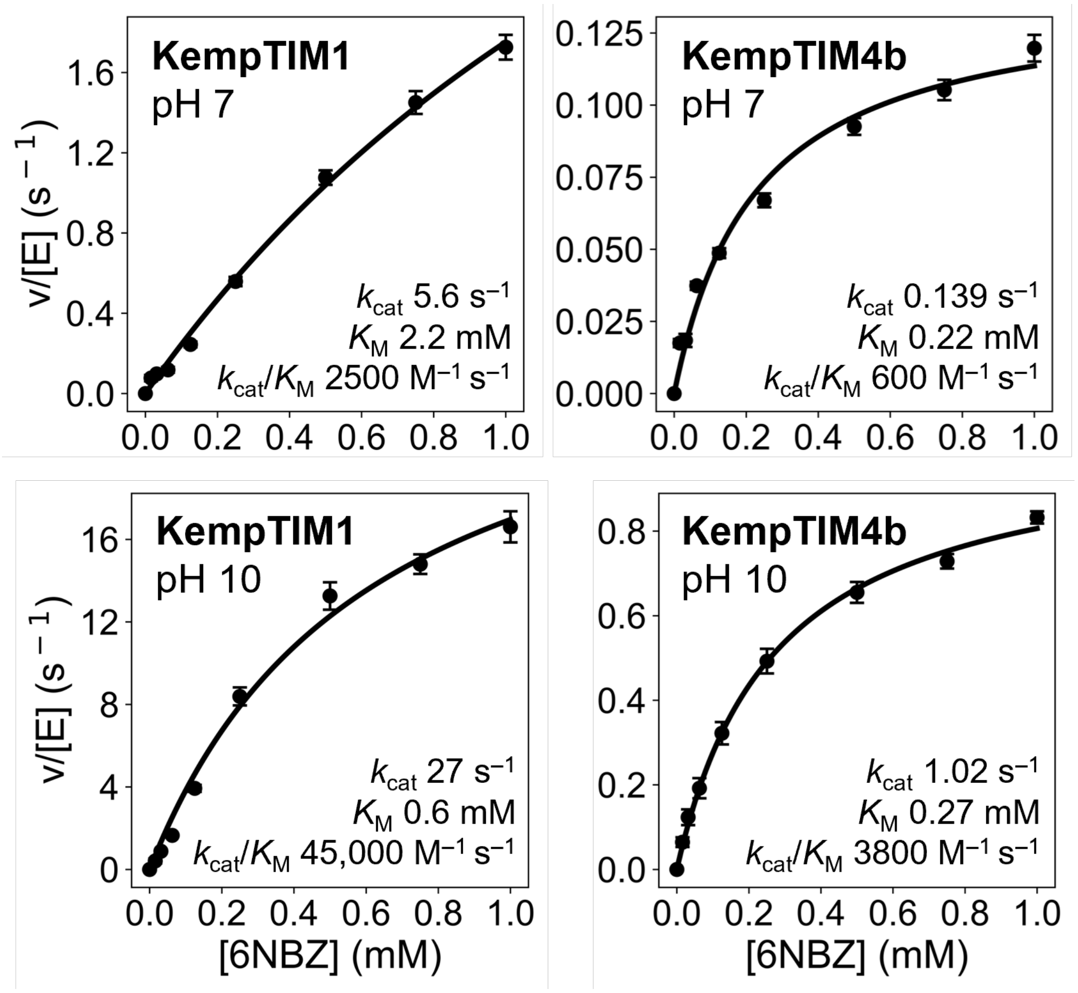


**Supplementary Figure 16. Kinetic characterization of KempTIMs with 6-nitrobenzisoxazole.** Michaelis-Menten plots showing normalized initial rates as a function of 6-nitrobenzisoxazole (6NBZ) concentration. Data represent the mean ± SEM of ≥ 6 replicate measurements from 2 independent protein batches. Kinetic assays were carried out in 50 mM sodium phosphate containing 100 mM NaCl and 10% MeOH (pH 7) or 50 mM CHES containing 100 mM NaCl and 10% MeOH (pH 10).

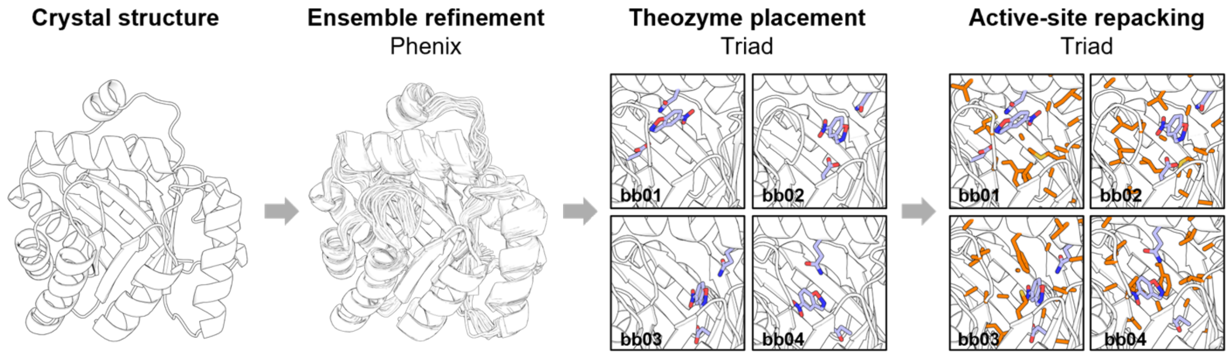


**Supplementary Figure 17. Ensemble-based enzyme design pipeline.** A conformational ensemble is generated from the crystal structure of a *de novo* enzyme using ensemble refinement in Phenix. Theozymes (blue sticks) containing an Asp catalytic base and either Asn or Gln as hydrogen-bond donors are then placed on each backbone template from the ensemble using the protein design software Triad. Distinct theozymes with catalytic residues at alternate positions are thus obtained for each backbone (bb01–bb04). Active sites (orange sticks) are subsequently repacked around the theozyme by optimizing the identity and side-chain conformations of residues within 4 Å of the transition state and catalytic residues, thereby maximizing favorable nonbonding interactions. Final designs are then filtered using structural metrics designed to reproduce mechanisms known to enhance enzyme catalysis.

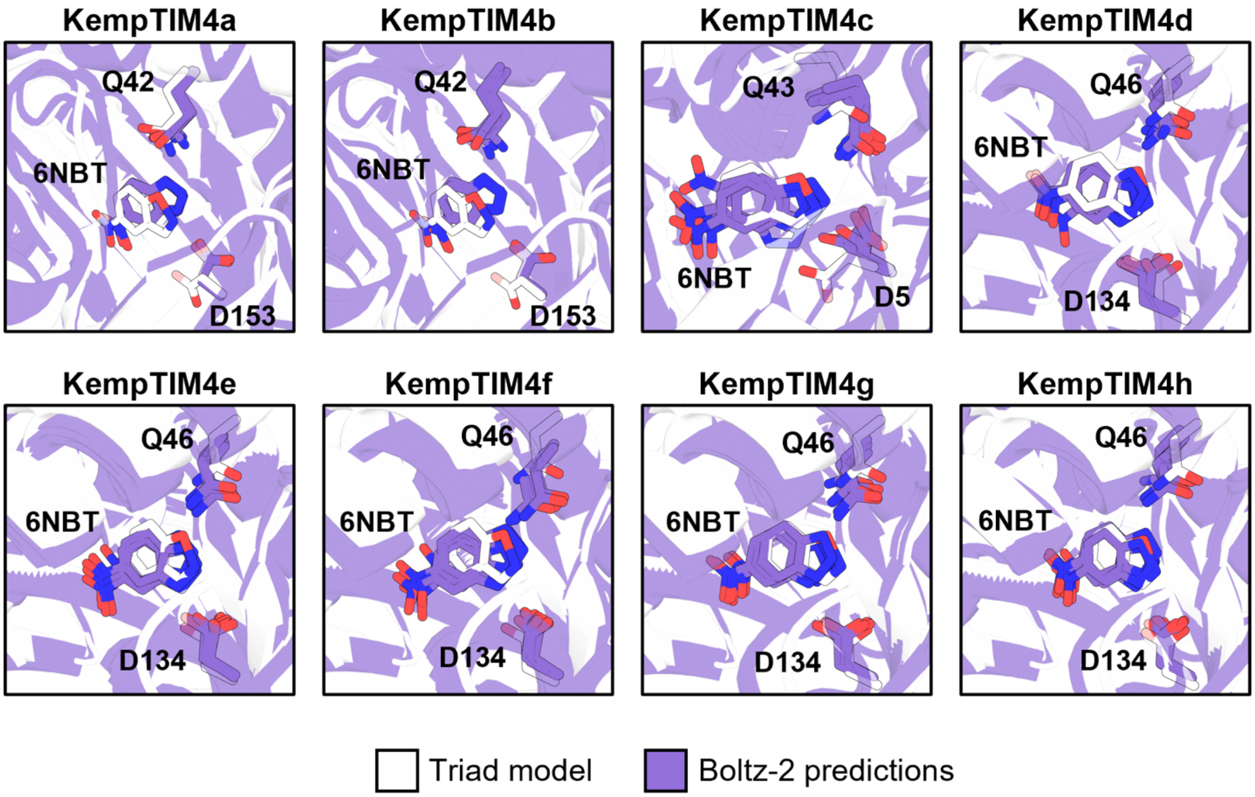


**Supplementary Figure 18. Boltz-2 predictions.** Five Boltz-2–predicted structures (purple) of KempTIM4 variants bound to the transition-state analogue 6-nitrobenzotriazole (6NBT) are overlaid on the Triad design model (white). The predicted 6NBT binding poses show strong agreement with the design model, with RMSDs < 1 Å in all cases. Catalytic contacts between designed residues and 6NBT are preserved, even when the catalytic residue adopts a different side-chain rotamer in the Boltz-2 models compared with the Triad model.

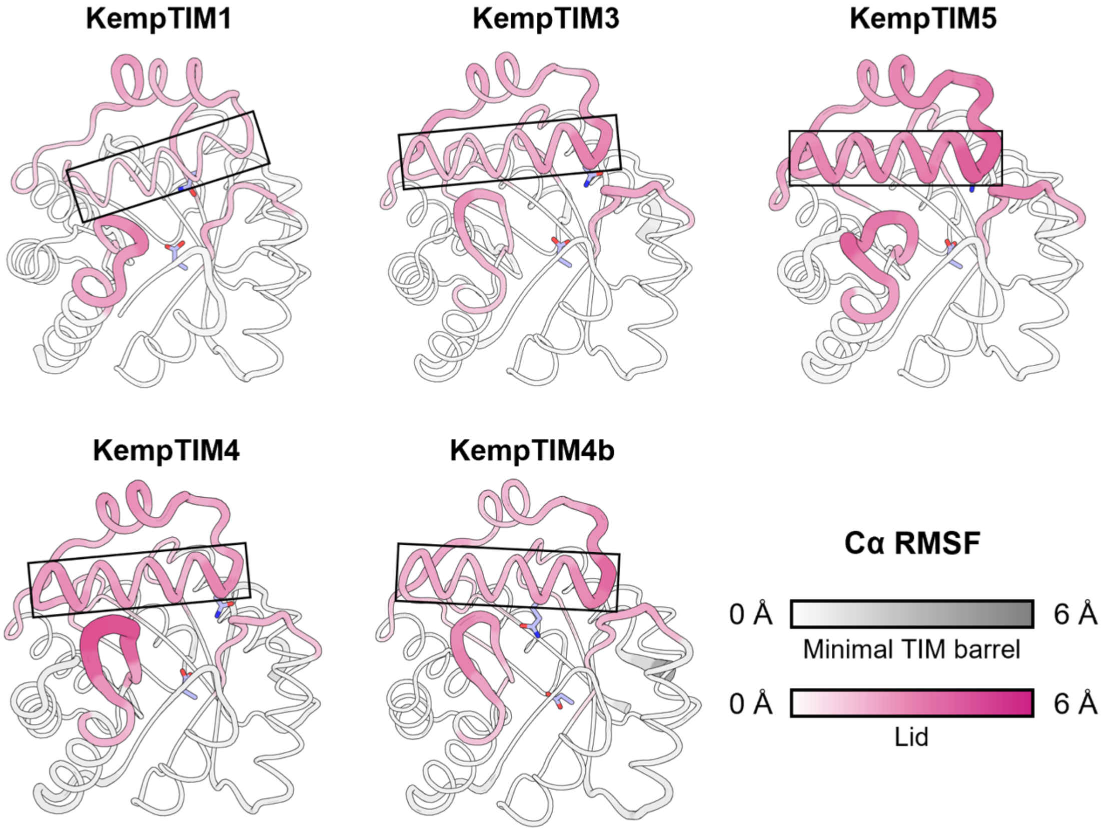


**Supplementary Figure 19. Molecular dynamics of KempTIMs.** Root-mean-square fluctuations (RMSF) of the protein backbone, shown as putty plots, indicate that the lids are consistently more flexible than the minimal TIM-barrel core across all designs. Within the lids, loop regions display the highest mobility, whereas the α-helix anchoring the designed catalytic H-bond donor (boxed) remains comparatively rigid. This helix is especially rigid in the most active first-round design, KempTIM1. Catalytic residues are shown as blue sticks.

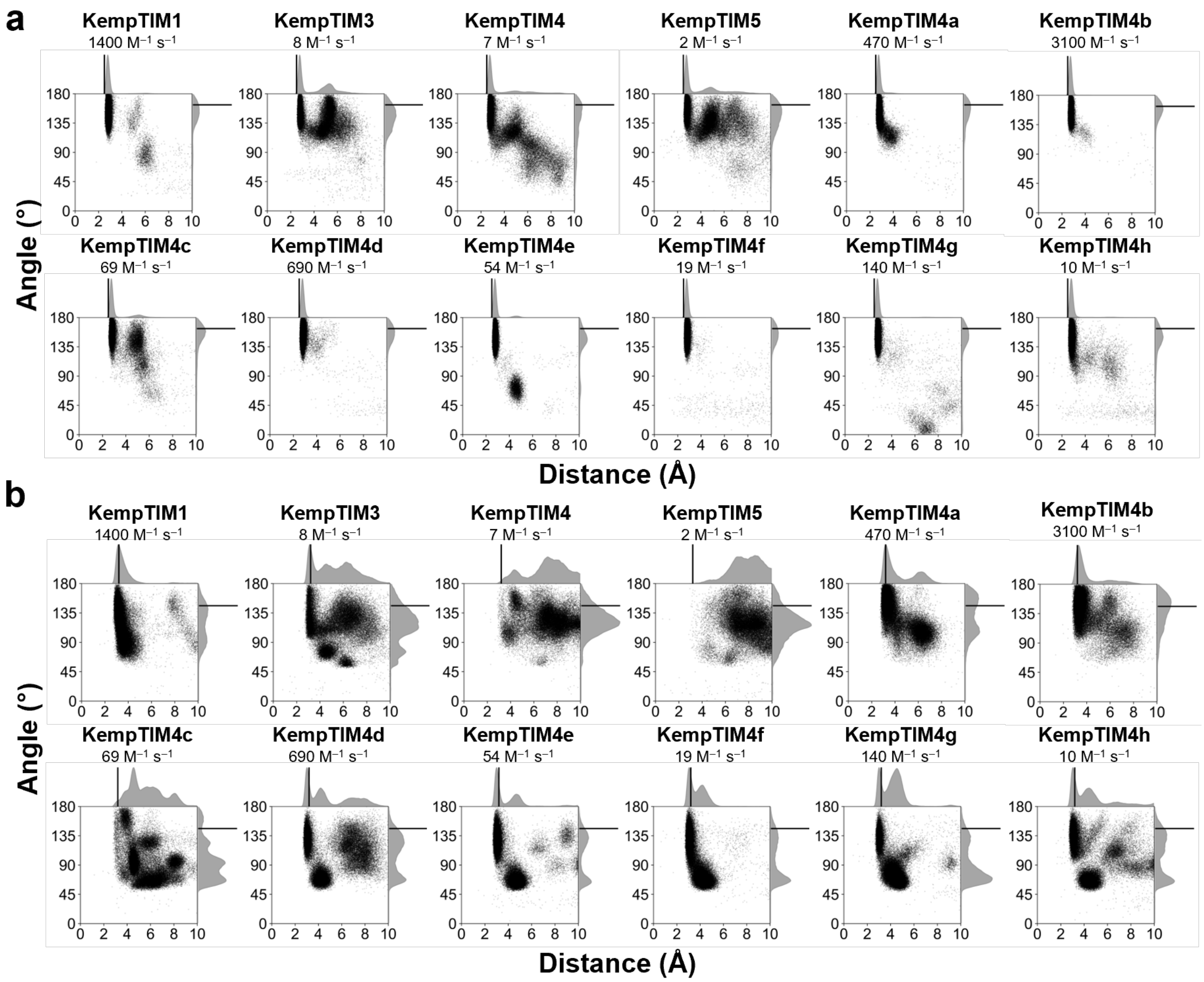


**Supplementary Figure 20. Catalytic contacts assessed by molecular dynamics (MD).** Scatter plots of angle versus distance for the base (top) and H-bond donor (bottom) contacts, compiled from ten 100-ns MD trajectories of KempTIM variants bound to the 6NBT transition-state analogue. For reference, the base/H-bond donor distances and angles in the crystal structure of HG3.17 (PDB ID: 5RGE)—2.5/3.2 Å and 163.0/146.1°, respectively— are shown as solid lines, representing ideal values associated with the high catalytic efficiency of this extensively evolved Kemp eliminase. Definitions of catalytic distances and angles between 6NBT and the designed catalytic residues are provided in Figure 5a. Catalytic efficiencies ($k_{cat}/K_M$) at pH 7 are listed for each variant.
